# Brain activity during facial processing in autism spectrum disorder: an activation likelihood estimation (ALE) meta-analysis of neuroimaging studies

**DOI:** 10.1101/2020.05.03.074351

**Authors:** Cristiano Costa, Ioana Alina Cristea, Elisa Dal Bò, Caterina Melloni, Claudio Gentili

**Affiliations:** Department of General Psychology, University of Padua, Italy; Department of Brain and Behavioral Sciences, University of Pavia, Pavia, Italy; Padova Neuroscience Center (PNC), University of Padua, Padua, Italy

**Keywords:** fmri, face perception, autism, ALE-Meta-analysis

## Abstract

**Background:** Though aberrant face processing is a hallmark of autistic spectrum disorder (ASD), findings on accompanying brain activity are divergent. Therefore, we conducted an activation likelihood estimation (ALE) meta-analysis of studies examining brain activity during face processing.

**Methods:** We searched PubMed and PsycINFO using combinations of terms as ‘fMRI’, ‘Autism Spectrum Disorder’, ‘Face Perception’. Eligible studies reported on DSM-diagnosed ASD patients, compared to controls (HC), using face stimuli presented in fMRI and reporting whole-brain analysis coordinates. We compared two approaches: “convergence of differences” (primary analysis) using study-level coordinates from ASD vs. HC contrasts, and “differences in convergence” (secondary) pooling coordinates within each group separately, and contrasting the resultant ALE-maps.

**Results:** Thirty-five studies (655 ASD and 668 HC) were included. Primary analysis identified a cluster in amygdala/parahippocampus where HC showed greater convergence of activation. Secondary analysis yielded no significant results.

**Conclusions:** Results suggest that ASD dysfunction in face processing relies on structures involved in emotional processing rather than perception. We also demonstrate that the two ALE methodologies lead to divergent results.

## 1. INTRODUCTON

Autism Spectrum Disorder (ASD) circumscribes a set of heterogeneous and lifelong neurodevelopmental disorders, defined by deficits in social communication and social interaction, and restricted, stereotyped and highly repetitive behaviours, interests or activities (*Diagnostic and statistical manual of mental disorders*◻*: DSM-5*, 2013).

Sensory deficits, already present in early developmental stages (Baranek et al., 2013), are cardinal characteristics of ASD and strong predictors of social communication and social interaction impairments (Turner-Brown et al., 2013), as well as of stereotyped and repetitive behaviour (Boyd et al., 2010). Specifically, ASD individuals show substantial deficits in face perception (Grelotti et al., 2002), owing to abnormal face processing strategies (Hobson et al., 1988), possibly caused by perceptual abnormalities, such as a locally oriented rather than global visual analysis (Morin et al., 2015), or more complex alterations of the social brain network (Pelphrey et al., 2014; Schultz et al., 2003). Impaired face perception could also underpin social interaction difficulties (Bi and Fang, 2017). Several studies (Dawson et al., 2005; Harms et al., 2010; Hileman et al., 2011) suggested that, compared to developmentally typical individuals, ASD patients show reduced accuracy and longer reaction times for identity or expression recognition.

Face perception is a highly sophisticated process subtended by two systems: the ‘core system’ and the ‘extended system’(Haxby et al., 2000). The ‘core system’ is mainly related to visual face processing. The ‘extended system’ includes non-visual areas extracting information from faces, such as the amygdala, insula, other limbic structures implicated in the emotional response to faces and other areas involved in autobiographic memory. Research on face perception in ASD suggested alterations in both systems, though findings were often inconsistent (Baron-Cohen et al., 2000; Robertson and Baron-Cohen, 2017). Abnormal brain activity in ASD individuals, specifically a reduced neural response, was identified in regions related to social cognition and face processing, such as the orbitofrontal cortex, superior temporal gyrus, amygdala (Baron-Cohen et al., 1999) and fusiform gyrus (Deffke et al., 2007). Yet despite a wealth of neuroimaging studies on sensory deficits in ASD, findings were inconsistent, revealing a multitude of abnormalities in early visual (Robertson and Baron-Cohen, 2017) or face-perception related areas (Weigelt et al., 2012), as well as in structures involved in emotional processing (Baron-Cohen et al., 2000).

Activation likelihood estimation (ALE) meta-analyses aim to summarize and identify consistency across neuroimaging findings. Briefly, this method computes the agreement of statistically significant foci across experiments in terms of probability distributions centered at the each set of focus coordinates (Eickhoff et al., 2009). Though it can only quantify convergence probabilities and not magnitude of activations, this method is particularly useful for fields with a suite of diverse and often inconsistent findings such as mental disorders, as it can theoretically parse out the most robust alterations in brain activity (Goodkind et al., 2015; Muller et al., 2017).

Two previous fMRI meta-analyses (Aoki et al., 2015; Nickl-Jockschat et al., 2015) examined emotional face processing in autism: one reported ASD-related hyperactivation in bilateral thalamus, caudate, and right precuneus, and ASD-related hypoactivation in the hypothalamus (Aoki et al., 2015). While, the other a cluster in the left fusiform gyrus due to reduce activations in ASD at single study level (Nickl-Jockschat et al., 2015). However, these meta-analysis used a small number of studies (13), including those relying on ROI analysis, a practice recently criticized (Eickhoff et al., 2016; Gentili et al., 2018; Müller et al., 2018).

Consequently, we conducted a systematic review and (ALE) meta-analysis of neuroimaging studies of face-related stimuli in individuals with ASD, with the aim of highlighting the more consistent neurobiological alterations. We also tested whether findings diverged depending on the two possible ALE meta-analysis approaches (Müller et al., 2018) (i.e., “differences in convergence” vs “convergence of differences”).

## 2 METHODS

### 2.1 Study selection

Eligible studies were identified by searching the National Library of Medicine/PubMed and PsycINFO bibliographic databases from inception until 4th of July 2019. We used combinations of database-specific terms as ‘fMRI’, ‘Autism’, ‘Face’, ‘Facial, ‘Visual Attention’, ‘Visual Processing’ ‘Fusiform Gyrus’ (figure 1 and Supplementary Material for the exact search string). Eligible studies were: (1) neuroimaging studies using functional magnetic resonance imaging (fMRI) in (2) participants of any age diagnosed with ASD according to DSM IV, IV-TR or 5, including comorbid disorders, (3) compared to a matched healthy control group (HC), (4) in a task employing faces or face parts (5) within the same experimental paradigm for both ASD and HC, (6) and conducting a direct univariate comparison of brain activation between ASD and HC (i.e., HC > ASD and/or ASD > HC), (7) for which 3D coordinates of peak activations in stereotactic space of the Montreal Neurological Institute (MNI) or Talairach were reported, (8) employing whole brain and not just to Region of Interest (ROI) analysis. Patients could be undergoing any kind of therapy (e.g., psychological, pharmacological). Reviews and meta-analyses were excluded. Two authors (CM, CG) independently screened and selected studies.

**Figure 1:**
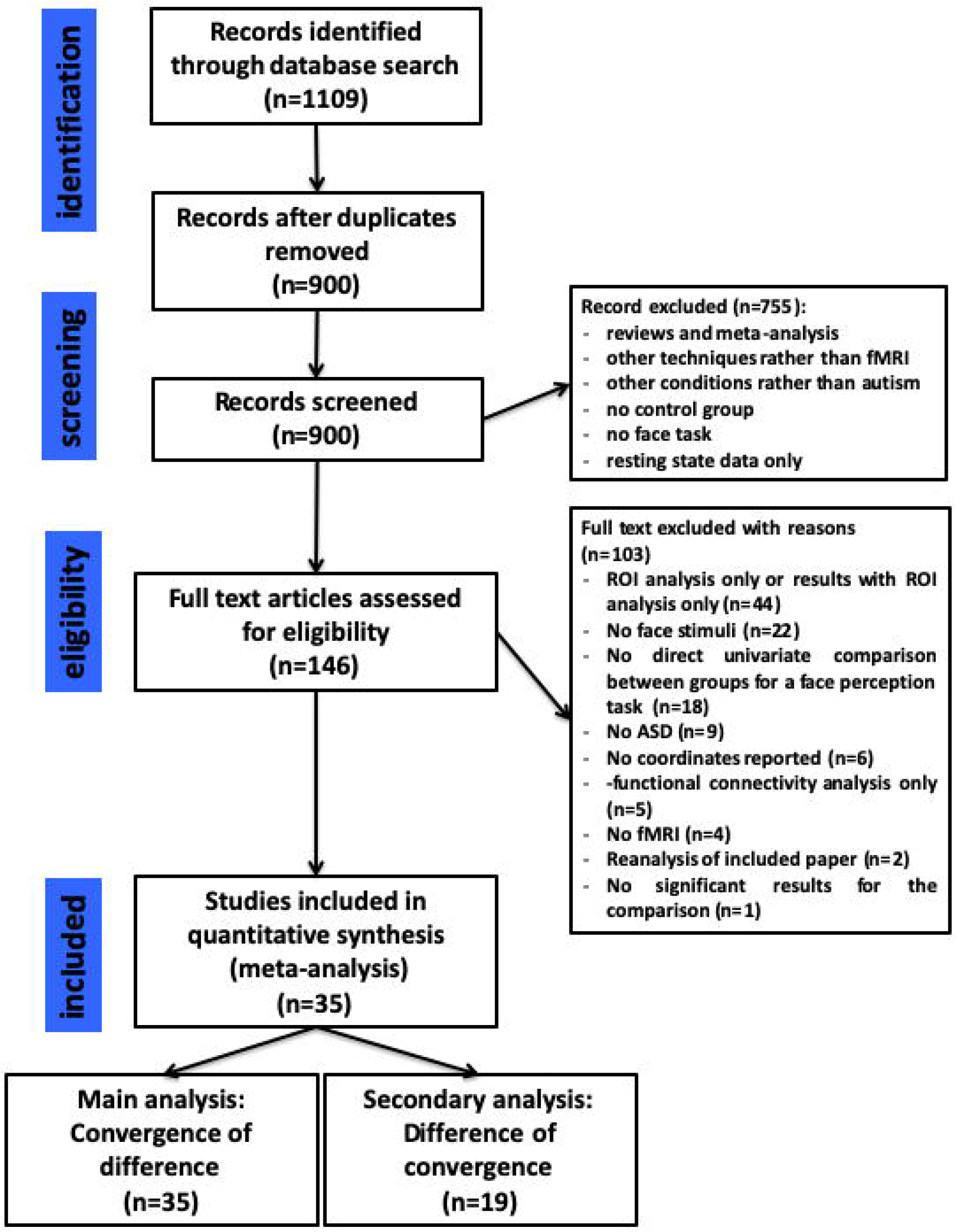
Prisma flow-chart illustrating the selection process of the present meta◻analysis

### 2.2 Data extraction

From each paper the following information as extracted: (1) participant gender and mean age; (2) diagnosis; (3) comorbidity; (4) concurrent treatments; (5) type of task and stimuli; (6) brain activation coordinates for the direct comparison between ASD and HC; (7) where available, activation coordinates within each single group (ASD and HC). Data were extracted independently by two researchers (CC, CM).

### 2.3 Study Quality

The quality and Risk of Bias (RoB) of included studies were evaluated with a modified version of the Newcastle-Ottawa scale (NOS)(Wells, 2001), (mNOS), adapted to fMRI data (Gentili et al., 2018). This version uses a different set of items adapted to fMRI studies (e.g., use of appropriate statistical corrections). Scores on the mNOS range from 0 to 11, with 0 to 3 considered indicative of high risk, 4 to 7 as intermediate and 8 to 11 as low risk. RoB was independently assessed by two researchers (CM, EDB). Inter-rater agreement was measured with the Kappa statistic, and disagreements were subsequently resolved by discussion with a third author (CG).

### 2.3 ALE meta-analysis

Stereotactic coordinates (x, y, z) were extracted from the studies, to be used in the Activation Likelihood Estimation (ALE) Meta-Analysis. The ALE algorithm was used as implemented in the GingerALE 2.3.6 software (Eickhoff et al., 2009). We used the correction for multiple comparisons derived from the same dataset implemented in GingerALE (Turkeltaub et al., 2012). Sample size for each foci experiment has been used to calculate the Full◻Width Half◻Maximum (FWHM) of the Gaussian function used to blur the foci. Coordinates in the MNI 152 standard space were converted into the Talairach space using the GingerALE foci converter tool.

Two approaches can be employed in an ALE meta-analysis of two groups. The first (“convergence of activation differences”) uses coordinates from the contrast ‘patients vs. controls’ (i.e., patients > controls and controls > patients). The second (“differences in convergence”) pools the activation reported within each group separately, and subsequently computes a contrast between the resultant ALE-maps. The two approaches have never been compared directly on the same data.

We used convergence of activation differences as the primary analysis because it used data from all included studies. We computed two independent meta-analysis (one for HC > ASD and the other for ASD > HC). Statistical significance was assessed and corrected for multiple comparisons using the cluster-wise method embedded in GingerALE: p < 0.001 cluster forming threshold, p < 0.01 cluster corrected FWE and N = 2,000 permutations.

To check the robustness of the findings, we also performed two sensitivity analyses. The first was a pooled analysis across ASD>HC and HC>ASD. This analysis might reflect a better summary of group differences as differences between analysis approaches and control conditions between single studies may have influenced the direction of group differences. Given the heterogeneity of tasks employed, we performed a second analysis limited to studies using solely face perception as task (see supplementary methods).

For the secondary analysis (differences in convergence), we computed a meta-analysis for activations of controls and ASD separately and contrasted them in a meta-analysis. For the single group meta-analysis, we used the same parameters described above, while to compute the differences of convergence, we used an uncorrected p value < 0.001, N=10000 permutations and a cluster threshold of 100 mm^3^. Gaussian smoothing for each meta-analysis was independently calculated by the software (Eickhoff et al., 2009).

This secondary analysis was restricted to studies that reported single group results, which were only a share of the entire pool. Therefore, differences between the primary and secondary analysis could be due to the different number of included studies and not to genuine divergences between the methods. To account for this possibility, we also conducted sensitivity analyses, in which the primary method (convergence of differences) was limited to the studies reporting the single group activations (Figure 1 and Supplementary Methods). To maintain consistency with the main analysis, we excluded one study (Zürcher et al., 2013) in which the contrast used in the single group analysis was different from that used in the convergence of difference. For each study we included coordinates for single groups analysis for the same contrasts used in the convergence of differences analysis or, if there was no such overlap, the most similar contrast (e.g. faces vs. baseline used in single group analysis and faces vs. objects and houses used in HC vs ASD analysis).

Finally, as post-hoc analysis, we examined whether results obtained with each of the two meta-analysis methods were also mirrored by the single studies. Specifically, for each included study, we checked whether (1) activation was reported in a cluster or region overlapping the one resulting from the meta-analysis and (2) if activation was present, whether it was discussed in the paper.

## 3 RESULTS

### 3.1 Study selection

The search produced 1109 entries (900 after removal of duplicates), 755 of which were excluded based on the abstract, i.e., failing to specify the method for diagnosing ASD or inadequately describing imaging methods. The remaining 146 were retrieved and full-texts were assessed. A total of 111 articles were excluded due to (1) lack of direct univariate comparison between ASD and HC (n=18), or comparison restricted to functional connectivity analysis (n=5) or no significant results for the comparison (n=1); (2) lack of reporting of coordinates for contrasts (n=6) or ROI only reported (n=44); and (3) re-analyses of previous, already included, studies (n=2); (4) lack of face stimuli in the task (n=22); (5) lack of fMRI use (n=4); (6) lack of ASD individual (e.g. use of autistic trait in HC) (n=9). A total of 35 articles (describing 36 experiments) were included in the meta-analysis, as described in the PRISMA flow diagram (figure 1).

### 3.2 Characteristics of included studies (Table 1, Table S1, Supplementary Results)

The 36 experiments included 1323 subjects (655 ASD and 668 HC). All studies performed whole-brain analyses: 17 reported both contrasts HC > ASD and ASD > HC, 15 the HC > ASD contrast only, whereas three the ASD > HC contrast only. Twenty-one studies also reported single group analyses (figure 1). Due to the limited number of studies including participants with comorbities or concomitant medication and to the reduced number of patients with these characteristics within these studies, we could not conduct further sensitivity analyses (Supplementary materials).

**Table 1.**
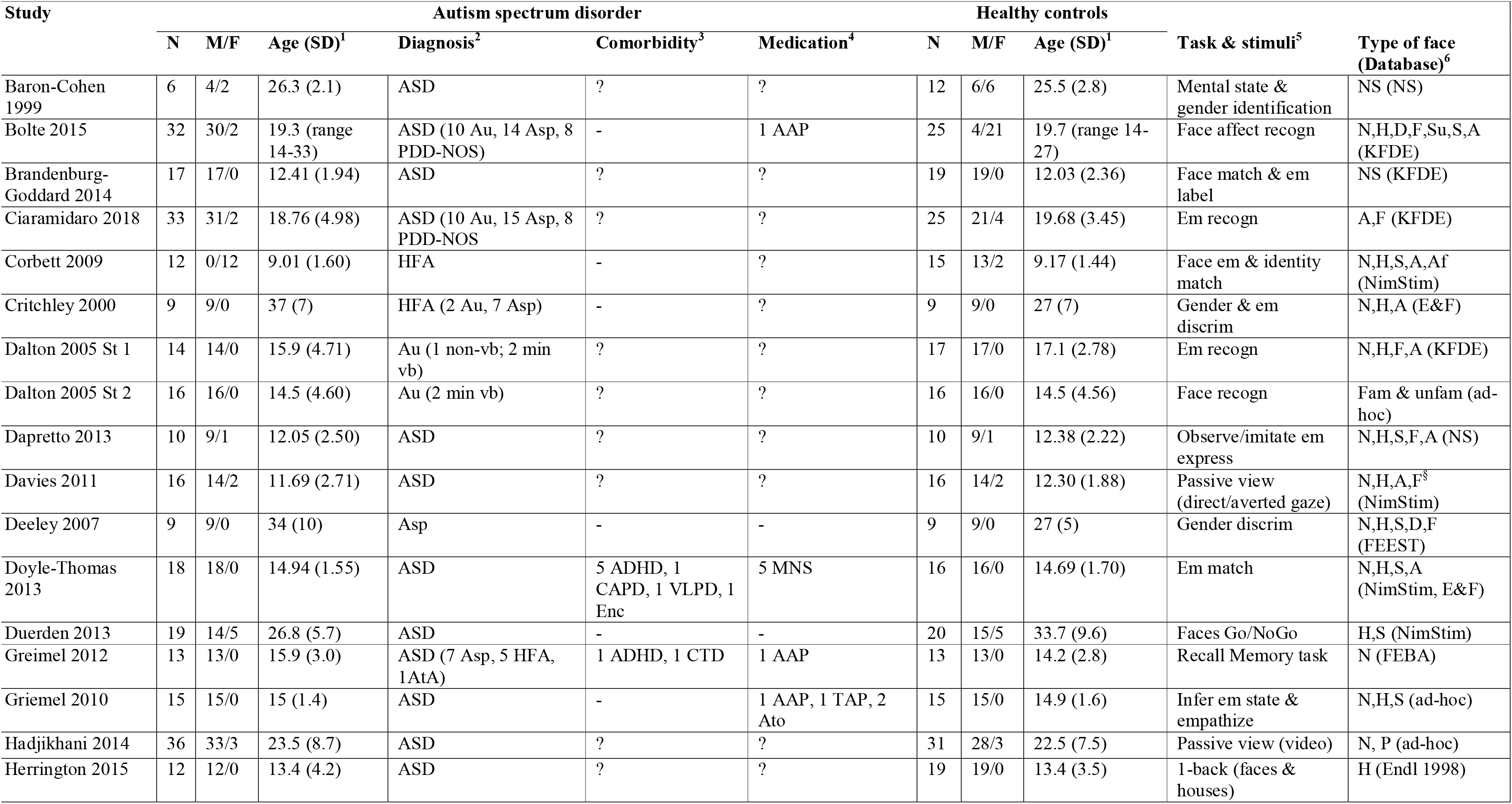
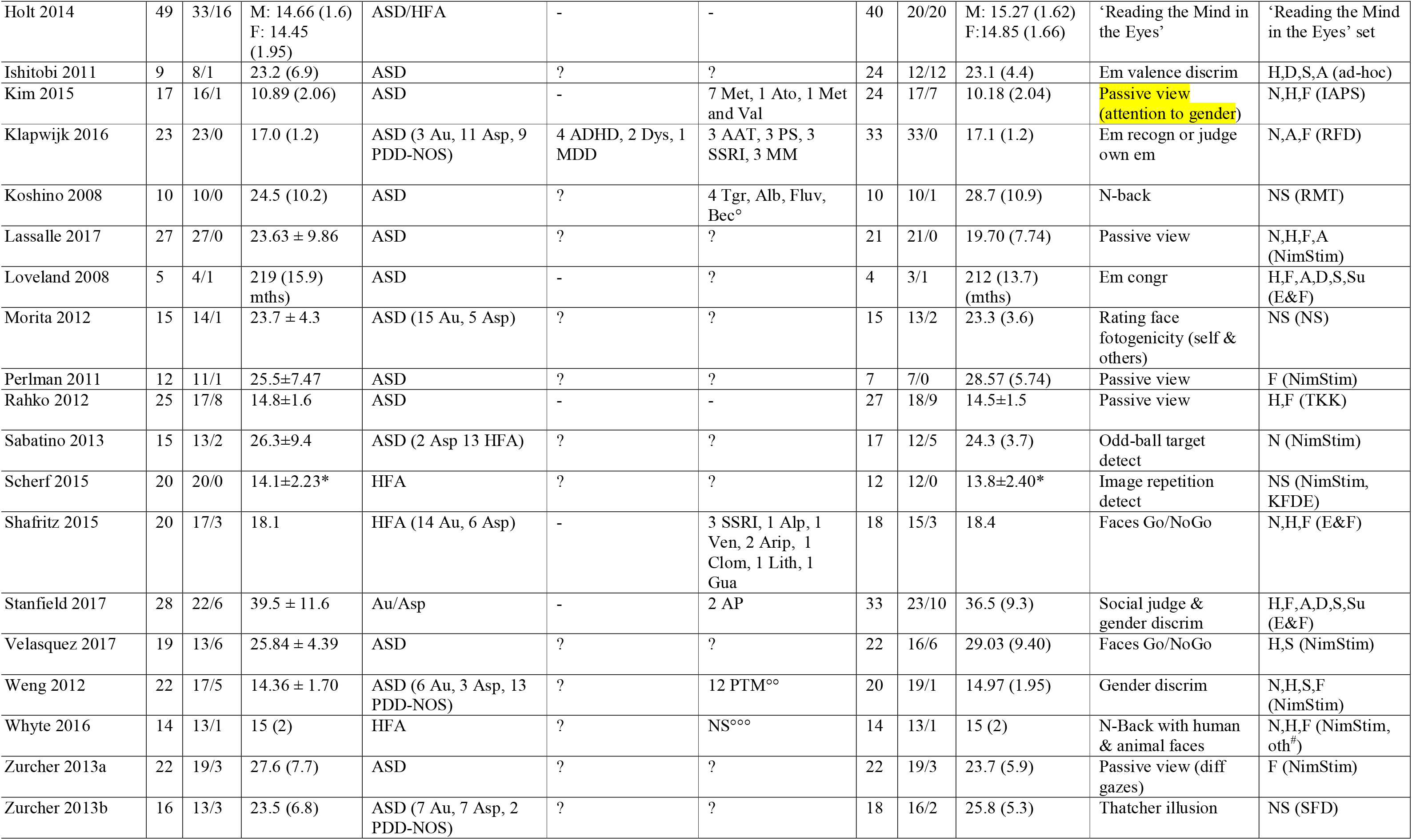
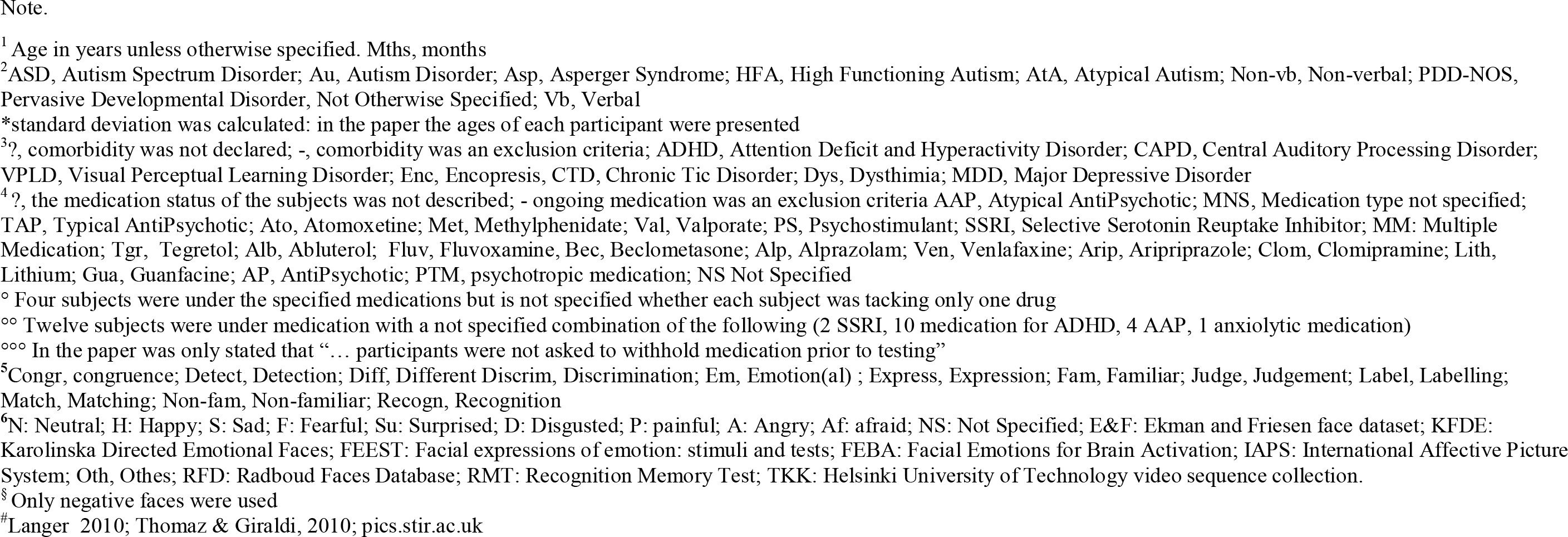
Characteristics of the studies included in the meta-analysis

| Study | Autism spectrum disorder |  |  |  |  |  | Healthy controls |  |  | Task & stimuli<sup>5</sup> | Type of face (Database)<sup>6</sup> |
| --- | --- | --- | --- | --- | --- | --- | --- | --- | --- | --- | --- |
|  | N | M/F | Age (SD)<sup>1</sup> | Diagnosis<sup>2</sup> | Comorbidity<sup>3</sup> | Medication<sup>4</sup> | N | M/F | Age (SD)<sup>1</sup> |  |  |
| Baron-Cohen 1999 | 6 | 4/2 | 26.3 (2.1) | ASD | ? | ? | 12 | 6/6 | 25.5 (2.8) | Mental state & gender identification | NS (NS) |
| Bolte 2015 | 32 | 30/2 | 19.3 (range 14-33) | ASD (10 Au, 14 Asp, 8 PDD-NOS) | - | 1 AAP | 25 | 4/21 | 19.7 (range 14-27) | Face affect recogn | N,H,D,F,Su,S,A (KFDE) |
| Brandenburg-Goddard 2014 | 17 | 17/0 | 12.41 (1.94) | ASD | ? | ? | 19 | 19/0 | 12.03 (2.36) | Face match & em label | NS (KFDE) |
| Ciaramidaro 2018 | 33 | 31/2 | 18.76 (4.98) | ASD (10 Au, 15 Asp, 8 PDD-NOS) | ? | ? | 25 | 21/4 | 19.68 (3.45) | Em recogn | A,F (KFDE) |
| Corbett 2009 | 12 | 0/12 | 9.01 (1.60) | HFA | - | ? | 15 | 13/2 | 9.17 (1.44) | Face em & identity match | N,H,S,A,Af (NimStim) |
| Critchley 2000 | 9 | 9/0 | 37 (7) | HFA (2 Au, 7 Asp) | - | ? | 9 | 9/0 | 27 (7) | Gender & em discrim | N,H,A (E&F) |
| Dalton 2005 St 1 | 14 | 14/0 | 15.9 (4.71) | Au (1 non-vb; 2 min vb) | ? | ? | 17 | 17/0 | 17.1 (2.78) | Em recogn | N,H,F,A (KFDE) |
| Dalton 2005 St 2 | 16 | 16/0 | 14.5 (4.60) | Au (2 min vb) | ? | ? | 16 | 16/0 | 14.5 (4.56) | Face recogn | Fam & unfam (ad-hoc) |
| Dapretto 2013 | 10 | 9/1 | 12.05 (2.50) | ASD | ? | ? | 10 | 9/1 | 12.38 (2.22) | Observe/imitate em express | N,H,S,F,A (NS) |
| Davies 2011 | 16 | 14/2 | 11.69 (2.71) | ASD | ? | ? | 16 | 14/2 | 12.30 (1.88) | Passive view (direct/averted gaze) | N,H,A,F<sup>§</sup> (NimStim) |
| Deeley 2007 | 9 | 9/0 | 34 (10) | Asp | - | - | 9 | 9/0 | 27 (5) | Gender discrim | N,H,S,D,F (FEEST) |
| Doyle-Thomas 2013 | 18 | 18/0 | 14.94 (1.55) | ASD | 5 ADHD, 1 CAPD, 1 VLDP, 1 Enc | 5 MNS | 16 | 16/0 | 14.69 (1.70) | Em match | N,H,S,A (NimStim, E&F) |
| Duerden 2013 | 19 | 14/5 | 26.8 (5.7) | ASD | - | - | 20 | 15/5 | 33.7 (9.6) | Faces Go/NoGo | H,S (NimStim) |
| Greimel 2012 | 13 | 13/0 | 15.9 (3.0) | ASD (7 Asp, 5 HFA, 1AtA) | 1 ADHD, 1 CTD | 1 AAP | 13 | 13/0 | 14.2 (2.8) | Recall Memory task | N (FEBA) |
| Griemel 2010 | 15 | 15/0 | 15 (1.4) | ASD | - | 1 AAP, 1 TAP, 2 Ato | 15 | 15/0 | 14.9 (1.6) | Infer em state & empathize | N,H,S (ad-hoc) |
| Hadjikhani 2014 | 36 | 33/3 | 23.5 (8.7) | ASD | ? | ? | 31 | 28/3 | 22.5 (7.5) | Passive view (video) | N, P (ad-hoc) |
| Herrington 2015 | 12 | 12/0 | 13.4 (4.2) | ASD | ? | ? | 19 | 19/0 | 13.4 (3.5) | 1-back (faces & houses) | H (Endl 1998) |
| Holt 2014 | 49 | 33/16 | M: 14.66 (1.6)<br>F: 14.45 (1.95) | ASD/HFA | - | - | 40 | 20/20 | M: 15.27 (1.62)<br>F: 14.85 (1.66) | 'Reading the Mind in the Eyes' | 'Reading the Mind in the Eyes' set |
| Ishitobi 2011 | 9 | 8/1 | 23.2 (6.9) | ASD | ? | ? | 24 | 12/12 | 23.1 (4.4) | Em valence discrim | H,D,S,A (ad-hoc) |
| Kim 2015 | 17 | 16/1 | 10.89 (2.06) | ASD | - | 7 Met, 1 Ato, 1 Met and Val | 24 | 17/7 | 10.18 (2.04) | Passive view (attention to gender) | N,H,F (IAPS) |
| Klapwijk 2016 | 23 | 23/0 | 17.0 (1.2) | ASD (3 Au, 11 Asp, 9 PDD-NOS) | 4 ADHD, 2 Dys, 1 MDD | 3 AAT, 3 PS, 3 SSRI, 3 MM | 33 | 33/0 | 17.1 (1.2) | Em recogn or judge own em | N,A,F (RFD) |
| Koshino 2008 | 10 | 10/0 | 24.5 (10.2) | ASD | ? | 4 Tgr, Alb, Fluv, Bec<sup>o</sup> | 10 | 10/1 | 28.7 (10.9) | N-back | NS (RMT) |
| Lassalle 2017 | 27 | 27/0 | 23.63 ± 9.86 | ASD | ? | ? | 21 | 21/0 | 19.70 (7.74) | Passive view | N,H,F,A (NimStim) |
| Loveland 2008 | 5 | 4/1 | 219 (15.9) mths) | ASD | - | ? | 4 | 3/1 | 212 (13.7) (mths) | Em congr | H,F,A,D,S,Su (E&F) |
| Morita 2012 | 15 | 14/1 | 23.7 ± 4.3 | ASD (15 Au, 5 Asp) | ? | ? | 15 | 13/2 | 23.3 (3.6) | Rating face fotogenicity (self & others) | NS (NS) |
| Perlman 2011 | 12 | 11/1 | 25.5±7.47 | ASD | ? | ? | 7 | 7/0 | 28.57 (5.74) | Passive view | F (NimStim) |
| Rahko 2012 | 25 | 17/8 | 14.8±1.6 | ASD | - | - | 27 | 18/9 | 14.5±1.5 | Passive view | H,F (TKK) |
| Sabatino 2013 | 15 | 13/2 | 26.3±9.4 | ASD (2 Asp 13 HFA) | ? | ? | 17 | 12/5 | 24.3 (3.7) | Odd-ball target detect | N (NimStim) |
| Scherf 2015 | 20 | 20/0 | 14.1±2.23* | HFA | ? | ? | 12 | 12/0 | 13.8±2.40* | Image repetition detect | NS (NimStim, KFDE) |
| Shafritz 2015 | 20 | 17/3 | 18.1 | HFA (14 Au, 6 Asp) | - | 3 SSRI, 1 Alp, 1 Ven, 2 Arip, 1 Clom, 1 Lith, 1 Gua | 18 | 15/3 | 18.4 | Faces Go/NoGo | N,H,F (E&F) |
| Stanfield 2017 | 28 | 22/6 | 39.5 ± 11.6 | Au/Asp | - | 2 AP | 33 | 23/10 | 36.5 (9.3) | Social judge & gender discrim | H,F,A,D,S,Su (E&F) |
| Velasquez 2017 | 19 | 13/6 | 25.84 ± 4.39 | ASD | ? | ? | 22 | 16/6 | 29.03 (9.40) | Faces Go/NoGo | H,S (NimStim) |
| Weng 2012 | 22 | 17/5 | 14.36 ± 1.70 | ASD (6 Au, 3 Asp, 13 PDD-NOS) | ? | 12 PTM<sup>oo</sup> | 20 | 19/1 | 14.97 (1.95) | Gender discrim | N,H,S,F (NimStim) |
| Whyte 2016 | 14 | 13/1 | 15 (2) | HFA | ? | NS<sup>ooo</sup> | 14 | 13/1 | 15 (2) | N-Back with human & animal faces | N,H,F (NimStim, oth<sup>#</sup>) |
| Zurcher 2013a | 22 | 19/3 | 27.6 (7.7) | ASD | ? | ? | 22 | 19/3 | 23.7 (5.9) | Passive view (diff gazes) | F (NimStim) |
| Zurcher 2013b | 16 | 13/3 | 23.5 (6.8) | ASD (7 Au, 7 Asp, 2 PDD-NOS) | ? | ? | 18 | 16/2 | 25.8 (5.3) | Thatcher illusion | NS (SFD) |

Note.
<sup>1</sup> Age in years unless otherwise specified. Mths, months
<sup>2</sup> ASD, Autism Spectrum Disorder; Au, Autism Disorder; Asp, Asperger Syndrome; HFA, High Functioning Autism; AtA, Atypical Autism; Non-vb, Non-verbal; PDD-NOS, Pervasive Developmental Disorder, Not Otherwise Specified; Vb, Verbal
\*standard deviation was calculated: in the paper the ages of each participant were presented
<sup>3</sup>?, comorbidity was not declared; -, comorbidity was an exclusion criteria; ADHD, Attention Deficit and Hyperactivity Disorder; CAPD, Central Auditory Processing Disorder; VPLD, Visual Perceptual Learning Disorder; Enc, Encopresis, CTD, Chronic Tic Disorder; Dys, Dysthymia; MDD, Major Depressive Disorder
<sup>4</sup>?, the medication status of the subjects was not described; - ongoing medication was an exclusion criteria AAP, Atypical AntiPsychotic; MNS, Medication type not specified;
TAP, Typical AntiPsychotic; Ato, Atomoxetine; Met, Methylphenidate; Val, Valporate; PS, Psychostimulant; SSRI, Selective Serotonin Reuptake Inhibitor; MM: Multiple Medication; Tgr, Tegretol; Alb, Abluterol; Fluv, Fluvoxamine, Bec, Beclometasone; Alp, Alprazolam; Ven, Venlafaxine; Arip, Aripiprazole; Clom, Clomipramine; Lith, Lithium; Gua, Guanfacine; AP, AntiPsychotic; PTM, psychotropic medication; NS Not Specified
<sup>o</sup> Four subjects were under the specified medications but is not specified whether each subject was taking only one drug
<sup>oo</sup> Twelve subjects were under medication with a not specified combination of the following (2 SSRI, 10 medication for ADHD, 4 AAP, 1 anxiolytic medication)
<sup>ooo</sup> In the paper was only stated that "... participants were not asked to withhold medication prior to testing"
<sup>5</sup> Congr, congruence; Detect, Detection; Diff, Different Discrim, Discrimination; Em, Emotion(al) ; Express, Expression; Fam, Familiar; Judge, Judgement; Label, Labelling; Match, Matching; Non-fam, Non-familiar; Recogn, Recognition
<sup>6</sup>N: Neutral; H: Happy; S: Sad; F: Fearful; Su: Surprised; D: Disgusted; P: painful; A: Angry; Af: afraid; NS: Not Specified; E&F: Ekman and Friesen face dataset; KFDE: Karolinska Directed Emotional Faces; FEEST: Facial expressions of emotion: stimuli and tests; FEBA: Facial Emotions for Brain Activation; IAPS: International Affective Picture System; Oth, Othes; RFD: Radboud Faces Database; RMT: Recognition Memory Test; TKK: Helsinki University of Technology video sequence collection.
<sup>§</sup> Only negative faces were used
<sup>#</sup>Langer 2010; Thomaz & Giraldi, 2010; pics.stir.ac.uk

### 3.2 Study quality (Supplementary Results, Figure S1, Table S2)

The overall Cohen kappa (Mean ±SD) was 0.88±0.12 ranging from 1 to 0.63. Consensus and Cohen kappa for each item of the mNOS were reported in Table S2 and Figure S1. The lower agreement was for definition (0.63) and selection (0.69) of controls. Three studies were considered as low RoB, twenty-nine as intermediate risk, and three as high risk of bias. A detailed description of the quality of each study is presented in the supplementary results.

### 3.3 Primary analysis: convergence of differences

For the voxel-wise whole-brain analysis all the 36 experiments were considered. For the HC > ASD meta-analysis we included 32 experiments and the simulation obtained a minimum cluster size was 920 mm³ while for the ASD > HC meta-analysis we included 20 experiments and a cluster size of 688 mm³. We identified a single significant cluster in which the difference for the contrast HC > ASD showed a significant convergence. The cluster mainly belonged to left amygdala (64.4%) extending to the parahippocampus (Table 2, Figure 2). Post-hoc analysis revealed that twelve studies reported amygdala activation for the contrast HC vs. ASD, comprising of a left-lateralized cluster in 6 and a bilateral cluster in 5, while only one paper reported a right-lateralized cluster.). Only one paper discussed the possible meaning of lateralization (Critchley et al., 2000).

**Table 2.** Significant clusters for the comparison between autism spectrum disorder (ASD) and healthy controls (HC) using the primary analysis convergence of difference method (*p*◻<◻0.01 corrected).

| Contrast | Hemi-<br>sphere | Region | Center of mass |  |  |  | Peak |  |  | Peak ALE<br>p value | Volume<br>(mm3) |
| --- | --- | --- | --- | --- | --- | --- | --- | --- | --- | --- | --- |
|  |  |  | BA | x | y | z | x | y | z |  |  |
| HC > ASD |  |  |  |  |  |  |  |  |  |  |  |
|  | L | Amygdala |  | -25.3 | -1.4 | -12.1 | -28 | -4 | -10 | 0.021 | 1112 |
|  | L | PHG | 34 |  |  |  | -24 | 0 | -14 | 0.020 |  |

*Note.* ASD: Autism Spectrum Disorder; HC: Healthy controls; L: left; R: right; PHG: Parahippocampal Gyrus

**Figure 2:**
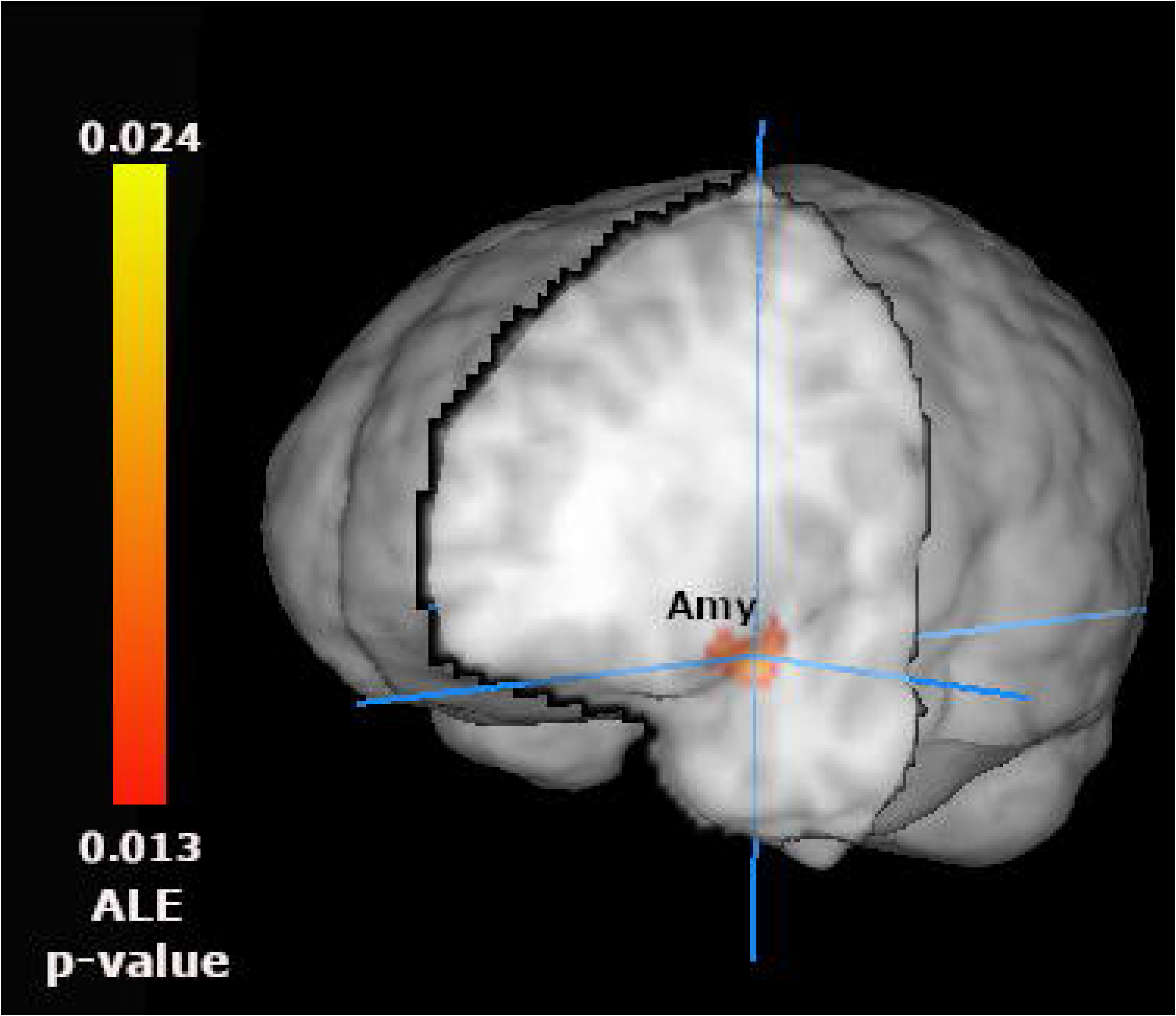
Significant results for the HC◻>◻ASD contrast of interest (*p*◻<◻0.01 corrected). Amy: amygdala. ALE p-value: Activation likelihood estimation probability

### 3.4 Secondary analysis: difference in convergences

A total of 21 experiments reported coordinates for single group analyses although two were excluded leading to 19 studies included in this analysis (19 for HC and 16 for ASD) (see Supplementary Methods and table S3). Results for the meta-analysis within each group are reported in the Supplement (Table S4, Figure S2). No significant clusters were identified for either (HC > ASD, ASD > HC) contrast.

### 3.5. Sensitivity analyses for the primary analysis

#### 3.5.1. ‘Pure’ face perception

The results were significant in the right amygdala for the HC > ASD (21 experiments) (Table S5 and Figure S3). For the ASD > HC meta-analysis (11 experiments) we did not find significant results.

#### 3.5.2. Pooled analysis

The pooled analysis found a convergence of differences in the two amygdalae (table S6 and Figure S4)

### 3.6. Sensitivity analyses for the secondary analysis

No significant cluster was evidenced for the ASD > HC meta-analysis (11 studies) using the same threshold of primary analysis. However, with a more liberal threshold (p <0.01 uncorrected) we found a significant cluster of convergence in a cluster including in the left amygdala and parahippocampus, largely overlapping with that in the primary analysis. (Table S7, Figure S5). The HC > ASD meta-analysis (18 experiments) did not yield significant results.

## 4 DISCUSSION

### 4.1 ALE meta-analysis results

In this voxel-wise whole-brain ALE meta-analysis, we did not uncover differences in convergence in the “core system” for face perception, particularly the fusiform gyrus, contradicting previous single studies (e.g.(Deffke et al., 2007)),. However, our findings support a crucial role for the “extended system”, confirmed the involvement of limbic and subcortical structures, such as the amygdala and parahippocampal gyrus. Specifically, in the primary analysis including all studies reporting direct comparisons between HC and ASD, we found differences in convergence in the left amygdala extended to the parahippocampal gyrus. Findings were supported in the pooled analysis, which revealed a bilateral amygdala cluster. Another sensitivity analysis limited to ‘pure’ face perception tasks also highlighted the altered activity of the amygdala, although with a different location (contralateral – right – amygdala). Examination of single studies indicated this difference was related to a higher activation of these regions in controls versus ASD patients, during visual processing of face stimuli.

The amygdala is crucial for emotional processing. Its abnormal activity may contribute to impairments in social interactions, face and emotional recognition (Donovan and Basson, 2017). Both structural and functional amygdala alterations were often reported in ASD patients (Donovan and Basson, 2017; Kemper and Bauman, 1993). For instance, adults with ASD showed no amygdala activation during the *‘Judging the Mind in the Eyes’* task, whereas healthy participants showed activation of the left amygdala (Baron-Cohen et al., 1999). In an in-depth examination of the included studies, we discovered that one third reported a unilateral amygdala activation, which was left localized in eight studies, and right localized in four. However, only one study included a discussion of lateralization (Baron-Cohen et al., 1999) (Table S8). Differences in convergence in the left amygdala lends further support to the oft-cited notion that the two amygdalae underpin different functions (Gainotti, 2018; Gläscher and Adolphs, 2003; Zalla et al., 2000), with the left involved in more “cold” cognitive and detailed processing of emotions (Dyck et al., 2011; Gainotti, 2018; Gläscher and Adolphs, 2003). As we included all studies involving faces as stimuli regardless of the task, our findings offer additional evidence for the specific involvement of the left amygdala in the ability of inferring mental state from complex visual stimuli (e.g., eyes region), frequently impaired in ASD (Baron-Cohen et al., 1999; Ketter et al., 1996).

Our data fails to replicate the results of two previous meta-analysis (Aoki et al., 2015; Nickl-Jockschat et al., 2015) which found ASD-related hyperactivation in thalamus, caudate, and precuneus, and ASD-related hypoactivation in the hypothalamus (Aoki et al., 2015) in one case and a ASD-related hypoactivation in the fusiform gyrus in the other (Nickl-Jockschat et al., 2015). We believe that the small number of studies included and the different inclusion criteria (e.g. using ROIs) account for most of the differences.

### 4.2 “Convergence of differences” OR “differences in convergence”?

From a methodological standpoint, we report on the first, to our knowledge, comparison within the same dataset of the two current ALE meta-analysis approaches: convergence of differences, which combines study-level activations for the contrast of ASD and HC, and, respectively, differences in convergence, which combines study-level activations within each group to compute two separate meta-analyses, one for ASD and one for HC, and subsequently contrasts these single group results (Müller et al., 2018). Such meta-analytic contrast highlighted the locations where in one group stronger convergence is found compared to the other.

We demonstrate that the two approaches yield highly divergent results. The first resulted into a significant cluster of convergence of differences in the left amygdala, whereas the second yielded no differences between groups. However, the second approach was limited to studies that reported results within single groups and consequently relied on fewer studies. To test for the possibility that divergences between the two methods would be explained by differences in the number of included studies, we conducted a sensitivity analysis applying the first method to the pool of studies used in the second. To test for the possibility that divergences between the two methods would be explained by differences in the number of included studies, we conducted a sensitivity analysis applying the first method to the pool of studies used in the second: a single cluster was evidenced, consistent with the primary findings.

Despite the limitation of this analysis (lower threshold and small number of experiments included – 11), it is unlikely that the divergent findings yielded by the two methods can be attributed to variations in the number of included studies. Rather, the discrepancy is probably grounded within the structure of ALE meta-analysis, which combines activations reported as significant within each study into a measure of convergence, i.e. declaring higher convergence if more studies reported activations in the same area. Unavoidably, the method draws heavily on the data analysis approach employed in each single study. For instance, a study with a more lenient or even inappropriate correction for the statistical threshold of activation will still contribute to convergence results. This problem is likely enhanced in meta-analyses examining convergence of single group activations (i.e., the 2^nd^ method) rather convergence of reported differences in activation. For instance, assuming an fMRI study uses 20 patients and 20 matched controls performing the same task, comparisons in brain activation between the two groups rely on more participants and therefore have more power than the examination of task-related activations within each group.

Moreover, examining convergence resulting from activations within single groups (e.g., patients or controls) rather than convergence resulting from contrasts between groups might obscure important differences, as well as elevate marginal ones. For instance, using the differences in convergence approach, we found no differences in the activation of the amygdala between ASD and controls, despite the fact one third of the studies reported significant activation for this contrast. This result is probably explained by the fact that the amygdala was activated, albeit differentially in the two groups resulting in a significant convergence within both ASD and HC. While difference in magnitude of activations are significant at a single experiment level in many cases, difference of convergence may not be significant.

### 4.3 Limitations and conclusions

One important limitation regards a considerable number of studies (n=73) that were excluded for not reporting between groups contrasts for the face perception task (n=18), performing only a comparison of functional connectivity (n=5), not providing brain activation coordinates for a contrast (n= 6), or reporting only ROI analyses (n=44). Since studies were not prospectively registered, the decision to not report or selectively report contrast data might have hinged on statistical significance, with negative or inconsistent findings suppressed. Although the final number of included studies are enough, some sensitivity analysis (e.g. those with less than 13 studies) is underpowered and needs to be considered as preliminary (Eickhoff et al., 2016). Furthermore, though all included studies used faces, tasks were heterogeneous and differences among them could account for the few significant findings reported in this meta-analysis. This is an unavoidable limitation of the ALE approach, which aims to highlight the commonalities across studies. However, given the limited number of experiments using ASD patients and faces, stricter inclusion criteria would have resulted into a restricted pool of studies and considerably reduced the power to reliably detect any differences. Another limit related to ALE approach as to every coordinate based meta-analysis is the risk of information lost as compared to maps meta-analysis (Salimi-Khorshidi et al., 2009): more differences between ASD and HC could be found in the original spatial maps and lost using this technique. However, coordinate-based meta-analysis represents a good trade-off given the relatively low amount of available original data. Finally, lack of pre-registration of the present meta-analysis is an important limitation.

Using ALE meta-analysis, we found support for a key role of amygdala dysfunctions in underpinning face processing in patients with autism spectrum disorders. Our findings would suggest that the core alteration of ASD relies on brain structures involved in emotional processing rather than perception, particularly since we did not report any significant differences in the core face perception system. Combining participant-level unthresholded maps from all eligible studies could offer a more definitive answer on brain activity alterations in ASD patients. Furthermore, we demonstrate that the two current ALE meta-analysis approaches can lead to highly divergent results. Neither represents a meta-analysis in a strict sense (Müller et al., 2018), since essential features such weighting of included studies or quantification of heterogeneity are absent (Higgins and Green, 2011) from the ALE methodology- or indeed any neuroimaging meta-analysis. Hence, both methods should be viewed as tools for descriptively summarizing neuroimaging literature. Crucially, only statistically significant results are combined in an ALE meta-analysis, leading to an unavoidably biased summary of the literature. These limitations notwithstanding, the approach based on the convergence of differences appears to mirror single study findings more closely and is thus probably better suited for summarizing available data. The more complex question as to whether either method describes ‘real’ rather than spurious differences in brain activity remains open.

## Author Contributions

Dr Gentili had full access to all the data in the study and takes responsibility for the integrity of the data and the accuracy of the data analysis.

*Study concept and design:* Gentili, Cristea

*Acquisition, analysis, or interpretation of data:* Costa, Dal Bò, Melloni, Gentili

*Statistical analysis:* Gentili, Costa, Melloni, Dal Bò

*Study supervision:* Gentili

*Manuscript Draft*: Costa, Gentili, Cristea

*Critical revision of the manuscript for important intellectual content*: Gentili, Cristea

*Review of the final version of the manuscript*: all

## Conflict of Interest Disclosures

None reported.

## Funder/Sponsor

None reported.

## Notes

### Competing Interest Statement

The authors have declared no competing interest.

## References

Aoki, Y., Cortese, S., Tansella, M., 2015. Neural bases of atypical emotional face processing in autism: A meta-analysis of fMRI studies. World J. Biol. Psychiatry Off. J. World Fed. Soc. Biol. Psychiatry 16, 291–300. https://doi.org/10.3109/15622975.2014.957719

Baranek, G.T., Watson, L.R., Boyd, B.A., Poe, M.D., David, F.J., McGuire, L., 2013. Hyporesponsiveness to social and nonsocial sensory stimuli in children with autism, children with developmental delays, and typically developing children. Dev. Psychopathol. 25, 307–320. https://doi.org/10.1017/S0954579412001071

Baron-Cohen, S., Ring, H.A., Bullmore, E.T., Wheelwright, S., Ashwin, C., Williams, S.C., 2000. The amygdala theory of autism. Neurosci. Biobehav. Rev. 24, 355–364.

Baron-Cohen, S., Ring, H.A., Wheelwright, S., Bullmore, E.T., Brammer, M.J., Simmons, A., Williams, S.C.R., 1999. Social intelligence in the normal and autistic brain: an fMRI study: *Social intelligence in the normal and autistic brain*. Eur. J. Neurosci. 11, 1891–1898. https://doi.org/10.1046/j.1460-9568.1999.00621.x

Bi, T., Fang, F., 2017. Impaired Face Perception in Individuals with Autism Spectrum Disorder: Insights on Diagnosis and Treatment. Neurosci. Bull. 33, 757–759. https://doi.org/10.1007/s12264-017-0187-1

Boyd, B.A., Baranek, G.T., Sideris, J., Poe, M.D., Watson, L.R., Patten, E., Miller, H., 2010. Sensory features and repetitive behaviors in children with autism and developmental delays. Autism Res. n/a-n/a. https://doi.org/10.1002/aur.124

Critchley, H.D., Daly, E.M., Bullmore, E.T., Williams, S.C.R., Van Amelsvoort, T., Robertson, D.M., Rowe, A., Phillips, M., McAlonan, G., Howlin, P., Murphy, D.G.M., 2000. The functional neuroanatomy of social behaviour. Brain 123, 2203–2212. https://doi.org/10.1093/brain/123.11.2203

Dawson, G., Webb, S.J., McPartland, J., 2005. Understanding the Nature of Face Processing Impairment in Autism: Insights From Behavioral and Electrophysiological Studies. Dev. Neuropsychol. 27, 403–424. https://doi.org/10.1207/s15326942dn2703_6

Deffke, I., Sander, T., Heidenreich, J., Sommer, W., Curio, G., Trahms, L., Lueschow, A., 2007. MEG/EEG sources of the 170-ms response to faces are co-localized in the fusiform gyrus. NeuroImage 35, 1495–1501. https://doi.org/10.1016/j.neuroimage.2007.01.034

Diagnostic and statistical manual of mental disorders◻: DSM-5, 2013. . Fifth edition. Arlington, VA◻: American Psychiatric Publishing, [2013] ©2013.

Donovan, A.P.A., Basson, M.A., 2017. The neuroanatomy of autism - a developmental perspective. J. Anat. 230, 4–15. https://doi.org/10.1111/joa.12542

Dyck, M., Loughead, J., Kellermann, T., Boers, F., Gur, R.C., Mathiak, K., 2011. Cognitive versus automatic mechanisms of mood induction differentially activate left and right amygdala. NeuroImage 54, 2503–2513. https://doi.org/10.1016/j.neuroimage.2010.10.013

Eickhoff, S.B., Laird, A.R., Grefkes, C., Wang, L.E., Zilles, K., Fox, P.T., 2009. Coordinate-based activation likelihood estimation meta-analysis of neuroimaging data: A random-effects approach based on empirical estimates of spatial uncertainty. Hum. Brain Mapp. 30, 2907–2926. https://doi.org/10.1002/hbm.20718

Eickhoff, S.B., Nichols, T.E., Laird, A.R., Hoffstaedter, F., Amunts, K., Fox, P.T., Bzdok, D., Eickhoff, C.R., 2016. Behavior, sensitivity, and power of activation likelihood estimation characterized by massive empirical simulation. NeuroImage 137, 70–85. https://doi.org/10.1016/j.neuroimage.2016.04.072

Gainotti, G., 2018. A historical review of investigations on laterality of emotions in the human brain. J. Hist. Neurosci. 1–19. https://doi.org/10.1080/0964704X.2018.1524683

Gentili, C., Messerotti Benvenuti, S., Lettieri, G., Costa, C., Cecchetti, L., 2018. ROI and phobias: The effect of ROI approach on an ALE meta◻analysis of specific phobias. Hum. Brain Mapp. https://doi.org/10.1002/hbm.24492

Gläscher, J., Adolphs, R., 2003. Processing of the Arousal of Subliminal and Supraliminal Emotional Stimuli by the Human Amygdala. J. Neurosci. 23, 10274–10282. https://doi.org/10.1523/JNEUROSCI.23-32-10274.2003

Goodkind, M., Eickhoff, S.B., Oathes, D.J., Jiang, Y., Chang, A., Jones-Hagata, L.B., Ortega, B.N., Zaiko, Y.V., Roach, E.L., Korgaonkar, M.S., Grieve, S.M., Galatzer-Levy, I., Fox, P.T., Etkin, A., 2015. Identification of a common neurobiological substrate for mental illness. JAMA Psychiatry 72, 305–315. https://doi.org/10.1001/jamapsychiatry.2014.2206

Grelotti, D.J., Gauthier, I., Schultz, R.T., 2002. Social interest and the development of cortical face specialization: What autism teaches us about face processing. Dev. Psychobiol. 40, 213–225. https://doi.org/10.1002/dev.10028

Harms, M.B., Martin, A., Wallace, G.L., 2010. Facial Emotion Recognition in Autism Spectrum Disorders: A Review of Behavioral and Neuroimaging Studies. Neuropsychol. Rev. 20, 290–322. https://doi.org/10.1007/s11065-010-9138-6

Haxby, J.V., Hoffman, E.A., Gobbini, M.I., 2000. The distributed human neural system for face perception. Trends Cogn. Sci. 4, 223–233. https://doi.org/10.1016/S1364-6613(00)01482-0

Higgins, J., Green, S. (Eds.), 2011. Cochrane Handbook for Systematic Reviews of Interventions Version 5.1.0 [updated March 2011]. The Cochrane Collaboration.

Hileman, C.M., Henderson, H., Mundy, P., Newell, L., Jaime, M., 2011. Developmental and Individual Differences on the P1 and N170 ERP Components in Children With and Without Autism. Dev. Neuropsychol. 36, 214–236. https://doi.org/10.1080/87565641.2010.549870

Hobson, R.P., Ouston, J., Lee, A., 1988. What’s in a face? The case of autism. Br. J. Psychol. 79, 441–453. https://doi.org/10.1111/j.2044-8295.1988.tb02745.x

Kemper, T.L., Bauman, M.L., 1993. The Contribution of Neuropathologic Studies to the Understanding of Autism. Neurol. Clin. 11, 175–187. https://doi.org/10.1016/S0733-8619(18)30176-2

Ketter, T.A., Andreason, P.J., George, M.S., Lee, C., Gill, D.S., Parekh, P.I., Willis, M.W., Herscovitch, P., Post, R.M., 1996. Anterior Paralimbic Mediation of Procaine-Induced Emotional and Psychosensory Experiences. Arch Gen Psychiatry 53, 59–69. https://doi.org/10.1001/archpsyc.1996.01830010061009

Morin, K., Guy, J., Habak, C., Wilson, H.R., Pagani, L., Mottron, L., Bertone, A., 2015. Atypical Face Perception in Autism: A Point of View? Autism Res. Off. J. Int. Soc. Autism Res. 8, 497–506. https://doi.org/10.1002/aur.1464

Müller, V.I., Cieslik, E.C., Laird, A.R., Fox, P.T., Radua, J., Mataix-Cols, D., Tench, C.R., Yarkoni, T., Nichols, T.E., Turkeltaub, P.E., Wager, T.D., Eickhoff, S.B., 2018. Ten simple rules for neuroimaging meta-analysis. Neurosci. Biobehav. Rev. 84, 151–161. https://doi.org/10.1016/j.neubiorev.2017.11.012

Muller, V.I., Cieslik, E.C., Serbanescu, I., Laird, A.R., Fox, P.T., Eickhoff, S.B., 2017. Altered Brain Activity in Unipolar Depression Revisited: Meta-analyses of Neuroimaging Studies. JAMA Psychiatry 74, 47–55. https://doi.org/10.1001/jamapsychiatry.2016.2783

Nickl-Jockschat, T., Rottschy, C., Thommes, J., Schneider, F., Laird, A.R., Fox, P.T., Eickhoff, S.B., 2015. Neural networks related to dysfunctional face processing in autism spectrum disorder. Brain Struct. Funct. 220, 2355–2371. https://doi.org/10.1007/s00429-014-0791-z

Pelphrey, K.A., Yang, D.Y.-J., McPartland, J.C., 2014. Building a social neuroscience of autism spectrum disorder. Curr. Top. Behav. Neurosci. 16, 215–233. https://doi.org/10.1007/7854_2013_253

Robertson, C.E., Baron-Cohen, S., 2017. Sensory perception in autism. Nat. Rev. Neurosci. 18, 671–684. https://doi.org/10.1038/nrn.2017.112

Salimi-Khorshidi, G., Smith, S.M., Keltner, J.R., Wager, T.D., Nichols, T.E., 2009. Meta-analysis of neuroimaging data: a comparison of image-based and coordinate-based pooling of studies. NeuroImage 45, 810–823. https://doi.org/10.1016/j.neuroimage.2008.12.039

Schultz, R.T., Grelotti, D.J., Klin, A., Kleinman, J., Van der Gaag, C., Marois, R., Skudlarski, P., 2003. The role of the fusiform face area in social cognition: implications for the pathobiology of autism. Philos. Trans. R. Soc. Lond. B. Biol. Sci. 358, 415–427. https://doi.org/10.1098/rstb.2002.1208

Turkeltaub, P.E., Eickhoff, S.B., Laird, A.R., Fox, M., Wiener, M., Fox, P., 2012. Minimizing within-experiment and within-group effects in activation likelihood estimation meta-analyses. Hum. Brain Mapp. 33, 1–13. https://doi.org/10.1002/hbm.21186

Turner-Brown, L.M., Baranek, G.T., Reznick, J.S., Watson, L.R., Crais, E.R., 2013. The First Year Inventory: a longitudinal follow-up of 12-month-old to 3-year-old children. Autism 17, 527–540. https://doi.org/10.1177/1362361312439633

Weigelt, S., Koldewyn, K., Kanwisher, N., 2012. Face identity recognition in autism spectrum disorders: A review of behavioral studies. Neurosci. Biobehav. Rev. 36, 1060–1084. https://doi.org/10.1016/j.neubiorev.2011.12.008

Wells, G., 2001. The Newcastle-Ottawa Scale (NOS) for assessing the quality of non randomised studies in meta-analyses. Httpwww Ohri Caprogramsclinicalepidemiologyoxford Asp.

Zalla, T., Koechlin, E., Pietrini, P., Basso, G., Aquino, P., Sirigu, A., Grafman, J., 2000. Differential amygdala responses to winning and losing: a functional magnetic resonance imaging study in humans: Amygdala response to winning and losing. Eur. J. Neurosci. 12, 1764–1770. https://doi.org/10.1046/j.1460-9568.2000.00064.x

Zürcher, N.R., Donnelly, N., Rogier, O., Russo, B., Hippolyte, L., Hadwin, J., Lemonnier, E., Hadjikhani, N., 2013. It’s All in the Eyes: Subcortical and Cortical Activation during Grotesqueness Perception in Autism. PLoS ONE 8, e54313. https://doi.org/10.1371/journal.pone.0054313

