## supplementary materials for "Brain activity during facial processing in autism spectrum disorder: an activation likelihood estimation (ALE) meta-analysis of neuroimaging studies"

**Supplementary content**

**Methods**

**Search string for National Library of Medicine/PubMed and PsychInfo**

The search was performed on the 4^th^ of July 2019 with the following search strings:

**PubMed**: ("magnetic resonance imaging"[MeSH Terms] OR ("magnetic"[All Fields] AND "resonance"[All Fields] AND "imaging"[All Fields]) OR "magnetic resonance imaging"[All Fields] OR "fmri"[All Fields]) AND ("autistic disorder"[MeSH Terms] OR ("autistic"[All Fields] AND "disorder"[All Fields]) OR "autistic disorder"[All Fields] OR "autism"[All Fields]) AND (("face"[MeSH Terms] OR "face"[All Fields]) OR ("face"[MeSH Terms] OR "face"[All Fields] OR "facial"[All Fields]) OR (visual[All Fields] AND ("attention"[MeSH Terms] OR "attention"[All Fields])) OR (visual[All Fields] AND processing[All Fields]) OR ("temporal lobe"[MeSH Terms] OR ("temporal"[All Fields] AND "lobe"[All Fields]) OR "temporal lobe"[All Fields] OR ("fusiform"[All Fields] AND "gyrus"[All Fields]) OR "fusiform gyrus"[All Fields])) resulting in 661 entries;

**PsychInfo**: (Autism OR ASD or autism spectrum disorder) AND (fmri OR functional magnetic resonance imaging OR brain imaging OR neuroimaging) AND (face OR faces OR facial OR visual attention OR visual perception OR visual processing OR fusiform gyrus) resulting in 448 entries.

**Study Quality:**

The modified version of the Newcastle-Ottawa scale (NOS) was described previously (1). The following domains from the original report were evaluated: (1) quality of sample selection based on i) case definition, ii) representativeness, iii) selection of the control group, and iv) definition of controls; (2) quality of reproducibility, i.e. whether the samples were comparable in terms of age, gender and other demographic variables considered in the study; 3) quality of exposure based on i) whether the two groups underwent the same experimental procedure (in our case this was an inclusion criteria), ii) whether drop-outs were described in the report, and iii) whether a difference in the behavioral task was shown. In the modified version, we introduced an additional 4th domain on the quality of data analysis, specific to fMRI: i) whether the studies used a sufficient cluster-forming threshold (i.e., voxel-based uncorrected p-value) to compensate for false positive underestimation, as recommended (2, 3)^,^ and ii) whether the authors used an appropriate statistical correction method (e.g. Clusterwise correction, voxelwise correction or False Discovery Rate correction) to avoid/reduce false positive results.

**Secondary analysis study inclusion:**

A total of 21 studies reported single group coordinates. Of these one was excluded because the coordinates were limited to ROI analysis and a second because the contrasts used for the single group analysis were different from those used in the contrasts between group (table S3).

**Sensitivity analyses:**

**Primary analysis: ‘pure’ face perception**

Two authors (CC, EDB) independently selected the studies based on the task. For this sensitivity analysis they included only studies in which a basic perception task was included (e.g., passive view, gender or emotional labelling) while studies including complex tasks (e.g. executive task such as go/no-go) where excluded. The discrepancies were solved through discussion and a Cohen Kappa was computed.

**Primary analysis: pooled analysis:**

A single meta-analysis for both HC > ASD and ASD > HC was computed using both the contrasts in the same gingerALE input. The statistical parameters were maintained identical to the primary analysis (p < 0.001 cluster forming threshold, p < 0.01 cluster corrected FWE and N = 2,000 permutations).

**Secondary analysis: convergence of difference**

We conducted a sensitivity analysis computing the convergence of difference including only those studies which present also group activations for ASD and HC separately. We used the same parameters of the primary and secondary analysis (p < 0.001 cluster forming threshold, p < 0.01 cluster corrected FWE and N = 2,000 permutations).

**Supplementary Results**

**Study selection and studies characteristics**

The 35 studies (5–39) included 1323 subjects (655 ASD and 668 HC). Only four papers report enrolling participants with a comorbid disorder (most frequently ADHD with a total of 13 cases in all the study considered). In all cases, the individuals with a comorbid disorder were under a half of the total patient group. Twelve studies listed comorbidity as exclusion criterion, while the remaining nine studies gave no details about comorbidities. Eleven studies included medicated patients (most frequently psychostimulants, antipsychotics and antidepressants). However, in all but one study, the medicated individuals totalled less than a half of the patient sample (two studies included only one medicated participant). Four studies listed concomitant medication as an exclusion criterion, while the remaining twenty made no mention of it.

**Study Quality**

Twenty-eight studies provided an adequate case definition, but in only eight, the sample could be considered representative of the population. Two studies provided an adequate selection of controls and three a satisfactory description of criteria used to select control participants. Regarding comparability, age and gender were matched between groups in thirty-two studies, whereas other variables (e.g., IQ) were controlled for in twenty-six. Fourteen studies reported dropout, and twenty-nine found differences in behavioural tasks. Regarding statistical quality, the adoption of a cluster-forming threshold to control for false positive (31, 32) results in neuroimaging studies (p<0.001) was used in only seven studies. Twenty-two studies reported the use of adequate methods to correct for multiple comparisons.

**Within group analysis**

The minimum cluster size was 840 mm³ for HC and 816 mm³ for ASD. Significant clusters for HC were located in two bilateral areas extended from the cerebellum to fusiform gyrus, and in the right amygdala (Table S4, Figure S2). Significant clusters for ASD were located in two bilateral areas extended from the cerebellum to fusiform gyrus (Table S4, Figure S2).

**Sensitivity analyses –**

**Primary analysis: “pure” face perception**

Two raters selected 24 studies as eligible, with an inter-rater agreement (Cohen Kappa) of 0.93 (the discrepancy was due to one study). Twenty-one experiments were used for the HC > ASD meta-analysis, and eleven for the ASD > HC meta-analysis. The HC > ASD meta-analysis yielded significant clusters (minimum volume: 616 mm^3^) in the amygdala region (Table S5, figure S3). The HC > ASD meta-analysis yielded no significant clusters of convergence. However, the number of studies included was small and the resultant small power could have precluded us from detecting differences.

**Primary analysis: pooled analysis**

Thirty-six experiments (in 35 studies) were used, resulting in three significant clusters of convergence (minimum volume: 624): in the two amygdalae and in the anterior cingulate cortex (Table S6, Figure S4).

**Secondary analysis: convergence of differences between groups in studies reporting single group activations**

Eighteen studies reported coordinates for HC > ASD contrast, and eleven for the ASD > HC contrast. Using the same parameters of the primary analysis (uncorrected p < 0.001, cluster-wise corrected p < 0.01), we did not detect any significant clusters. However, for a more liberal uncorrected threshold (uncorrected p < 0.01), we found a minimum cluster size of 2656 mm³ and a large (3616 mm^3^) cluster of significant convergence, including 9 peaks, which is overlapping the result of the primary analysis (Figure S5, Table S7). The ASD > HC meta-analysis did not yield significant clusters (minimum volume: 1600 mm^3^), even at the more liberal threshold, a finding that might also be explained by limited power due to the small number of studies.

**Supplementary Tables**

**Table S1. Description of contrasts/experiments (exp) reporting a direct comparison between autistic spectrum disorder patients and healthy controls in each included study**

| **Study** | **Multiple exp meta-analysis** | **Contrast/**  **Exp (performed)** | **Contrast/**  **Exp (meta-analysis)** | **Coordinates source** | **Notes** |
| --- | --- | --- | --- | --- | --- |
| Baron-Cohen 1999 | - | Theory of mind task | NA | Table 2 |  |
| Bolte 2015 | +(-) | Expl FAR vs. obj recogn  Impl FAR vs. obj recogn  Pre vs. post training | Expl FAR vs. obj recogn  Impl FAR vs. obj recogn | Table 2 |  |
| Brandenburg-Goddard 2014 | +(-) | Fa match > obj match  Fa label > Fa match | Fa label > Fa match | Table 3 | Only Fa label > Fa match was significant for HC vs. ASD |
| Ciaramidaro 2018 | + | Expl FAR vs Obj recogn  Impl FAR vs Obj recogn  N FAR vs Obj recogn | NA | Table 3 |  |
| Corbett 2009 | +(-) | Em > Ctrl Match Task  Person > Ctrl Match Task | Em > Ctrl Match Task | Table 4 | Only Em >Ctrl Match Task was significant for HC vs. ASD |
| Critchley 2000 | +(-) | Main effect of task:  Impl vs. expl em proc  Main effect of group:  HC vs. ASD  Group x task interaction | Main effect of group | Table 2 |  |
| Dalton 2005 Study 1 | - | Fa perc | NA | Figure 3 |  |
| Dalton 2005 Study 2 | - | Fa perc | NA | Figure 4 |  |
| Dapretto 2013 | +(-) | Fa perc  Fa imitation | Fa imitation | Table s3 | Only Fa Imitation Task was significant for HC vs. ASD |
| Davies 2011 | + | Direct - Null  Averted - Null  Direct - Averted | NA | Table 3 |  |
| Deleey 2007 | +(-) | F vs. bsln  H vs. bsln  D vs. bsln  Linear trend of act for em intensity | F vs. bsln  H vs. bsln  D vs. bsln | Table 3 |  |
| Doyle-Thomas 2013 | +(-) | Visual, Audio and Audiovisual em match | Audiovisual em match | Table 4 | Only Audiovisual em match was significant for HC vs. ASD |
| Duerden 2013 | - | NoGo vs. Go | NA | Table 4 |  |
| Greimel 2010 | +(-) | Other vs. high lvl bsln  Self vs. high lvl bsln  Father of ASD vs. Father of HC  ASD vs. HC | Other vs. high lvl bsln  For ASD vs. HC | In the text |  |
| Greimel 2012 | +(-) | Recol vs. NonRecol  RecolFa vs. NonRecolFa  RecolHouse vs. NonRecolHouse | RecolFace vs. NonRecolFace | In the text |  |
| Hadjikhani 2014 | - | Pain > No pain | NA | Table 3 |  |
| Herrington 2015 | - | Attend Fa > Attend House | NA | Table 2 |  |
| Holt 2014 | +(-) | Reading the Mind in the Eyes  HC vs. ASD  HC vs. Siblings | HC vs. ASD | Table 2 | Contrasts including siblings were not considered |
| Ishitobi 2011 | +(-) | Whole Fa vs. Bsln  Upper part of Fa vs. Bsln  Lower part of Fa vs. Bsln | Whole Fa vs. Bsln | Table 3 | No significant clusters for Upper part of Fa vs. Bsln and  Lower part of Fa vs. Baseline in HC vs. ASD |
| Kim 2015 | - | Fa passive viewing | NA | Table 2 |  |
| Klapwijk 2016 | +(-) | Em recogn > self-judgment on em  Contrast (em resonance)  HC vs. ASD  HC vs. CC/CU  CC/CU vs. ASD | HC vs. ASD | Table 2 | Contrasts including CC/CU were not considered |
| Koshino 2008 | - | N-Back Task | NA | Table 4 |  |
| Lassalle 2017 | + | ASD > HC for F 40 vs. N  HC > ASD for H 100 vs. N  ASD > HC for A 100 vs. A 40  HC > ASD for H 100 vs. H 40  HC > ASD for F 100 vs. F 40 | NA | Table 2  Table 3  sTable 10  sTable 11  sTable 12 |  |
| Loveland 2008 | - | Em vs. gender discrimination | NA | Table 3 |  |
| Morita 2012 | - | Self vs. Others | NA | Table 3 |  |
| Pearlman 2011 | +(-) | Free passive view vs. Bsln  Three diff types of Fa fixation | Free passive view vs. Bsln | Figure 2 | The contrast HC vs. ASD for the three fixation conditions were only performed with ROI analysis |
| Rahko 2012 | - | Passive view | NA | Table 1 |  |
| Sabatino 2013 | +(-) | Fa targets  HAI targets  Group x target interaction | Fa targets | Table 2 |  |
| Scherf 2015 | - | Passive view | NA | Table 3 |  |
| Shafritz 2015 | +(-) | ‘X’ NoGo vs. letter Go  H Go vs. F Go  F NoGo vs. H NoGo  H NoGo vs. F NoGo  Em NoGo vs. letter ‘X’ Nogo | H Go vs. F Go  F NoGo vs. H NoGo  H NoGo vs. F NoGo  Em NoGo vs. letter ‘X’ NoGo | Table 4 |  |
| Stanfield 2017 | +(-) | Main effect of task:  Social vs. gender judgment  Main effect of group:  HC vs. SPD vs. ASD  Group x condition interaction | Group x condition interaction btwn ASD and HC | Table s7 |  |
| Velasquez 2017 | +(-) | Letter NoGo > Go  Fa NoGo > Go  Fa NoGo > Letter NoGo | Fa NoGo > Go  Fa NoGo > Letter NoGo | Table 3 |  |
| Weng 2012 | + | Fa perc with gender discrimination | NA | sTable 4 |  |
| Whyte 2016 | +(-) | Human Fa vs. Obj  Animal Fa vs. Obj | Human Fa vs. Obj | Table 2 | No significant clusters for:  HC vs ASD for Animal > Object |
| Zurcher 2013a | - | Averted vs. Direct gaze | NA | Table 3 |  |
| Zurcher 2013b | - | Eyes up vs. Eyes inv | NA | Table 3 |  |

*Notes*: “+”, multiple contrasts/experiments reported; “-“, single contrasts/experiments reported; “+(-)”, multiple contrasts/experiments reported, but only one included in the meta-analysis. “Vs.”, Versus generically indicates the comparison between groups or condition without specify the sign (e.g., HC vs. ASD means both HC > ASD and ASD > HC). Exp, Experiment; Obs, Observation. FAR: Facial affect recognition; ASD: Autism Spectrum Disorder; HC: Healthy Controls; SPD: Schizotypal Personality Disorder; KS: Klinefelter syndrome; Expl, explicit; Impl, implicit; Fa, Face; Obj, object; Recogn, recognition; Match, matching; Label, labelling; N, neutral; F, Fearfull, H, Happy; A, Angry; D, Disgusted; act, activation; Em, emotion(al); Ctrl, control; Recol, recollected; NonRecol, non-recollected Proc, processing; Bsln, baseline; Lvl; level; View, viewing; Diff, different; HAI: High Autistic Interest; Btwn, between; perc, perception; Inv, inverted.

**Table S2. Study-level risk of bias ratings (after consensus) and the kappa inter-rater agreement (before consensus)**

|  | **Selection** | | | | **Comparability** | | **Exposure** | | | **Statistical analysis** | |
| --- | --- | --- | --- | --- | --- | --- | --- | --- | --- | --- | --- |
|  | **Study** | **Case def** | **Sel ctrl** | **Def ctrl** | **Age, sex** | **Oth** | **Exp** | **Drop** | **Behav** | **Clust** | **FP** |
| Baron-Cohen 1999 | ? | ? | ? | - | + | + | + | - | + | - | ? |
| Bolte 2015 | + | + | ? | ? | + | + | + | - | + | ? | + |
| Brandenburg-Goddard 2014 | + | - | - | ? | + | + | + | - | + | - | + |
| Ciaramidaro 2018 | + | ? | ? | ? | + | + | + | - | + | ? | + |
| Corbett 2009 | + | - | - | - | + | - | + | - | + | + | + |
| Critchley 2000 | + | - | ? | ? | - | + | + | - | + | + | ? |
| Dalton 2005 | + | - | ? | - | + | + | + | - | + | + | ? |
| Dapretto 2013 | ? | ? | ? | - | + | + | + | - | + | - | - |
| Davies 2011 | + | ? | ? | - | + | ? | + | - | + | - | + |
| Deeley 2007 | + | - | + | ? | - | + | + | **-** | - | - | + |
| Doyle-Thomas 2013 | + | - | - | - | + | + | + | ? | - | ? | + |
| Duerden 2013 | + | ? | ? | ? | - | - | + | - | - | ? | + |
| Greimel 2012 | + | - | ? | ? | + | + | + | - | + | + | ? |
| Griemel 2010 | + | - | ? | - | + | + | + | - | + | + | + |
| Hadjikhani 2014 | ? | - | - | - | + | + | + | + | + | ? | ? |
| Herrington 2015 | + | ? | ? | ? | + | ? | + | + | + | - | + |
| Holt 2014 | + | + | + | ? | + | + | + | + | + | ? | + |
| Ishitobi 2011 | + | + | - | ? | + | - | + | + | - | + | ? |
| Kim 2015 | + | - | - | + | + | + | + | + | + | - | - |
| Klapwijk 2016 | + | - | - | ? | + | + | + | - | + | - | + |
| Koshino 2008 | ? | - | ? | ? | + | - | + | - | + | ? | ? |
| Lassalle 2017 | + | - | - | ? | + | - | + | + | + | - | + |
| Loveland 2008 | + | - | ? | ? | + | + | + | - | + | - | + |
| Morita 2012 | + | + | - | ? | + | + | + | - | + | - | ? |
| Perlman 2011 | ? | ? | ? | ? | + | - | + | - | ? | ? | + |
| Rahko 2012 | + | - | - | + | + | + | + | + | + | - | + |
| Sabatino 2013 | - | + | - | - | + | + | + | - | + | - | + |
| Scherf 2015 | ? | ? | ? | - | + | ? | + | + | + | - | + |
| Shafritz 2015 | + | + | - | ? | + | + | + | + | + | - | ? |
| Stanfield 2017 | + | + | - | + | + | + | + | - | + | - | ? |
| Velasquez 2017 | + | ? | ? | - | + | + | + | + | + | - | + |
| Weng 2012 | + | + | ? | - | + | + | + | + | - | - | ? |
| Whyte 2016 | + | ? | ? | ? | + | + | + | + | + | - | + |
| Zurcher 2013a | + | - | - | ? | + | + | + | + | + | - | + |
| Zurcher 2013b | + | ? | ? | - | + | + | + | + | + | + | + |
| **Cohen’s Kappa** | **1.0** | **0.94** | **0.69** | **0.63** | **0.92** | **1.0** | **1.0** | **0.94** | **0.84** | **1.0** | **0.78** |

*Note.* Beh, behavioral; Clust, cluster forming threshold; Ctrl, control group; Def, definition; Diff, difference; Exp, exposure; FP, False-positives correction; Repres, representativeness;

**Table S3. Studies and contrasts within studies included in the secondary analysis**

| **Study** | **ASD** | **HC** | **ASD>HC** | **HC>ASD** |
| --- | --- | --- | --- | --- |
| Baron-Cohen 1999 | Theory of Mind task | Theory of Mind task | Theory of Mind task | Theory of Mind task |
| Brandenburg-Goddard 2014 |  | Face matching > Object matching; Face labeling > Face matching | Face matching > Object matching; Face labeling > Face matching |  |
| Ciaramidaro 2018 | Implicit FAR vs Object recognition | Implicit FAR vs Object recognition |  | Implicit FAR vs Object recognition |
| Corbett 2009 | Emotion recognition > Control; identity recognition > Control | Emotion recognition > Control; identity recognition > Control | Emotion recognition > Control; identity recognition > Control | Emotion recognition > Control; identity recognition > Control |
| Dapretto 2013 | Imitation of emotional expression | Imitation of emotional expression |  | Imitation of emotional expression |
| Davies 2011 | Direct - Null; Averted - Null | Direct - Null; Averted - Null | Direct - Null; Averted - Null | Direct - Null;  Averted - Null |
| Deleey 2007 | Fear, Happy | Fear, Disgust, Happy |  | Fear, Disgust, Happy, Sad |
| Doyle-Thomas 2013 | Audiovisual > Visual, Visual > Audiovisual, Audiovisual > Auditory, Auditory > Audiovisual | Audiovisual > Visual, Visual > Audiovisual, Audiovisual > Auditory, Auditory > Audiovisual | Audiovisual | Audiovisual |
| Duerden 2013 | NoGo vs. Go | NoGo vs. Go | NoGo vs. Go | NoGo vs. Go |
| Hadjikhani 2014 | Pain > No Pain | Pain > No Pain | Pain > No Pain | Pain > No Pain |
| Ishitobi 2011 | Whole face condition, Upper face condition, Lower face condition | Whole face condition, Upper face condition, Lower face condition |  | Whole face condition, Upper face condition, Lower face condition |
| Koshino 2008 | Working memory task | Working memory task | Working memory task | Working memory task |
| Lassalle 2017* | Fearful 100 vs Fearful 40  Fearful 40 vs Neutral | Happy 100 vs Happy 40,  Fearful 40 vs Neutral  Happy 100 vs Neutral | Fearful 40 vs Neutral | Happy 100 vs Neutral, Fearful 100 vs Fearful 40, Happy 100 vs Happy 40 |
| Loveland 2008 |  | Emotional > Sex |  | Emotional > Sex |
| Morita 2012 | Self vs Other | Self vs Other |  | Self vs Other |
| Scherf 2015 | Faces, Houses, Objects | Faces, Houses, Objects |  | Faces > Other (Objects+Houses) |
| Shafritz 2015 | Happy go > fear go  Emotional No-Go > Letter No-Go | Happy no-go vs fear no-go  Emotional No-Go > Letter No-Go | Happy no-go vs fear no-go  Emotional No-Go > Letter No-Go | Happy go > fear go |
| Stanfield 2017 | Social > Gender, Gender > Social | Social > Gender, Gender > Social |  | Social > Gender, Gender > Social |
| Zurcher 2013a |  | Averted > Direct |  | Averted > Direct |
| Tot = 19 | Tot = 16 | Tot = 19 | Tot = 9 | Tot = 18 |

* of note one of the contrasts of this paper (Fearful100 vs. Fearful40) has to be excluded because it is present in the HC vs. ASD comparisons but the supplementary table (sTable 9) which should have included the contrast for HC and ASD was listed but not published.

**Table S4 Significant clusters of convergence during facial processing within each of the HC and ASD groups (p < 0.01 corrected)**

|  |  |  |  | **Center of Mass** | | | **Peak** | | |  |  |
| --- | --- | --- | --- | --- | --- | --- | --- | --- | --- | --- | --- |
| **Group** | **Hemi-sphere** | **Region** | **BA** | **x** | **y** | **z** | **x** | **y** | **z** | **ALE** | **Volume** |
| **HC** |  |  |  |  |  |  |  |  |  |  |  |
|  | R | Culmen |  | 40 | -49.4 | -13.1 | 38 | -46 | -20 | 0.033 | 4096 |
|  | R | MTG | 21 |  |  |  | 52 | -44 | 10 | 0.019 |  |
|  | R | Fgyr | 37 |  |  |  | 42 | -62 | -6 | 0.017 |  |
|  | L | MTG | 37 |  |  |  | 48 | -52 | 0 | 0.016 |  |
|  | L | Fgyr | 37 | -43 | -62.3 | -10.3 | -44 | -64 | -10 | 0.026 | 2008 |
|  | R | Amygdala |  | 17.8 | -6.1 | -9.9 | 18 | -6 | -10 | 0.042 | 1696 |
|  | R | Fgyr | 19 | 32.1 | -80.4 | -12.3 | 28 | -84 | -12 | 0.019 | 880 |
|  | R | Fgyr | 19 |  |  |  | 36 | -78 | -12 | 0.018 |  |
| **ASD** |  |  |  |  |  |  |  |  |  |  |  |
|  | R | Culmen |  | 35.5 | -54.7 | -14.2 | 34 | -54 | -16 | 0.025 | 4040 |
|  | R | Declive |  |  |  |  | 38 | -44 | -20 | 0.023 |  |
|  | R | FGyr | 19 |  |  |  | 34 | -70 | -10 | 0.021 |  |
|  | R | MTG |  |  |  |  | 38 | -58 | -4 | 0.015 |  |
|  | L | Culmen |  | -40 | -47.6 | -18.9 | -40 | -46 | -20 | 0.027 | 1768 |

*Note.* HC, Healthy controls; ASD, Autism Spectrum Disorder; L, left; R, right; MTG, Middle Temporal Gyrus; Fgyr, Fusiform Gyrus. Each cluster is separated by a horizontal line: a single cluster may have multiple peaks.

**Table S5: Sensitivity analysis: pure face perception. Significant clusters for the comparison between autism spectrum disorder (ASD) and healthy controls (HC) using the primary analysis convergence of difference method (p < 0.01 corrected).**

|  |  |  |  |  | **Center of mass** | | **Peak** | | |  |
| --- | --- | --- | --- | --- | --- | --- | --- | --- | --- | --- |
| **Contrast** | **Hemi-sphere** | **Region** | **BA** | **x** | **y** | **z** | **x** | **y** | **z** | **Volume** |
| **HC >ASD** |  |  |  |  |  |  |  |  |  |  |
|  | R | PHG | 34 | 18.6 | -7.2 | -16 | 18 | -8 | -16 | 944 |

*Note.* HC, Healthy controls; ASD, Autism Spectrum Disorder; L, left; R, right; PHG, Parahippocampal Gyrus.

**Table S6: Sensitivity analysis: pooled analysis. Significant clusters considering coordinates for the ASD > HC and for the HC > ASD pooled together (p < 0.01 corrected).**

| **Hemi-sphere** | **Region** | **BA** | **x** | **y** | **z** | **x** | **y** | **z** | **Peak ALE p value** | **Volume** |
| --- | --- | --- | --- | --- | --- | --- | --- | --- | --- | --- |
| L | PHG | 34 | -23.9 | -4.2 | -12.7 | -24 | 0 | -12 | 0.028 | 1728 |
| L | Amygdala |  |  |  |  | -22 | -8 | -16 | 0.024 |  |
| L | Amygdala |  |  |  |  | -22 | -8 | -10 | 0.022 |  |
| R | Amygdala |  | 19.5 | -4.8 | -16 | 20 | -6 | -20 | 0.029 | 1248 |
| R | Putamen |  |  |  |  | 20 | 4 | -10 | 0.019 |  |

*Note.* HC, Healthy controls; ASD, Autism Spectrum Disorder; L, left; R, right; PHG, Parahippocampal Gyrus. Each cluster is separated by a horizontal line: a single cluster may have multiple peaks.

**Table S7. Sensitivity analysis: secondary analysis. Significant clusters for the comparison between autism spectrum disorder (ASD) and healthy controls (HC) in studies reporting single group activation using the convergence of difference method (p < 0.01 corrected).**

|  |  |  |  |  | **Center of mass** | | **Peak** | | |  |  |
| --- | --- | --- | --- | --- | --- | --- | --- | --- | --- | --- | --- |
| **Contrast** | **Hemi-sphere** | **Region** | **BA** | **x** | **y** | **z** | **x** | **y** | **z** | **Peak ALE p-value** | **Volume** |
| **HC > ASD** |  |  |  |  |  |  |  |  |  |  |  |
|  | L | SCG | 34 | -28.8 | 1.6 | -8.6 | -24 | 8 | -10 | 0.016 | 3616 |
|  | L | Insula | 13 |  |  |  | -42 | 10 | -4 | 0.015 |  |
|  | L | Amygdala |  |  |  |  | -22 | -8 | -8 | 0.013 |  |
|  | L |  | 13 |  |  |  | -36 | 6 | -8 | 0.013 |  |
|  | L | Putamen |  |  |  |  | -28 | -2 | -8 | 0.011 |  |
|  | L | PHG |  |  |  |  | -18 | -12 | -16 | 0.011 |  |
|  | L | Putamen |  |  |  |  | -22 | -2 | 0 | 0.010 |  |
|  | L | STS | 38 |  |  |  | -48 | 12 | -10 | 0.009 |  |
|  | L | Amygdala |  |  |  |  | -30 | -8 | -14 | 0.009 |  |

*Note.* HC, Healthy controls; ASD, Autism Spectrum Disorder; L, left; R, right; SCG, Subcallosas Gyrus; PHG, Parahippocampal Gyrus; STS, Superior Temporal Sulcus

**Table S8. Post-hoc study-level examination of significant clusters evidenced with the convergence of differences (Amygdala).**

|  | **Amygdala** | | | |
| --- | --- | --- | --- | --- |
| **Study** | **Reported** | **Contrast** | **Lateralized** | **Lateralization Discussed** |
| Baron-Cohen 1999 | + | HC>ASD | Left | + |
| Bolte 2015 | + | HC>ASD | Bilateral | N/A |
| Brandenburg-Goddard 2014 | + | ASD>HC | Right | - |
| Ciaramidaro 2018 | + | HC>ASD | Bilateral | N/A |
| Corbett 2009 | - | N/A | N/A | N/A |
| Critchley 2000 | + | HC>ASD | Left | + |
| Dalton 2005 Study 1 | + | ASD>HC | Left | - |
| Dalton 2005 Study 2 | + | ASD>HC | Right | - |
| Dapretto 2013 | + | ASD>HC | Bilateral | N/A |
| Davies 2011 | - | N/A | N/A | N/A |
| Deleey 2007 | - | N/A | N/A | N/A |
| Doyle-Thomas 2013 | - | N/A | N/A | N/A |
| Duerden 2013 | - | N/A | N/A | N/A |
| Greimel 2010 | - | N/A | N/A | N/A |
| Greimel 2012 | - | N/A | N/A | N/A |
| Hadjikhani 2014 | - | N/A | N/A | N/A |
| Herrington 2015 | - | N/A | N/A | N/A |
| Holt 2014 | - | N/A | N/A | N/A |
| Ishitobi 2011 | + | HC>ASD | Bilateral | N/A |
| Kim 2015 | + | HC>ASD | Right | - |
| Klapwijk 2016 | + | HC>ASD | Left | - |
| Koshino 2008 | - | N/A | N/A | N/A |
| Lassalle 2017 | + | ASD>HC | Left | - |
|  |  | HC>ASD | Bilateral | N/A |
| Loveland 2008 | - | N/A | N/A | N/A |
| Morita 2012 | - | N/A | N/A | N/A |
| Pearlman 2011 | + | HC>ASD | Bilateral | N/A |
| Rahko 2012 | - | N/A | N/A | N/A |
| Sabatino 2013 | + | HC>ASD | Left | - |
| Scherf 2015 | - | N/A | N/A | N/A |
| Shafritz 2015 | + | ASD>HC | Left | - |
|  |  | HC>ASD | Left | - |
| Stanfield 2017 | - | N/A | N/A | - |
| Velasquez 2017 | - | N/A | N/A | N/A |
| Weng 2012 | - | N/A | N/A | N/A |
| Whyte 2016 | + | HC>ASD | Left | - |
| Zurcher 2013a | - | N/A | N/A | N/A |
| Zurcher 2013b | - | N/A | N/A | N/A |

*Note.* “-“, absent, “+” present, N/A, not applicable

**Supplementary Figures**

**Figure S1: Study quality according to the modified version of the Newcastle‐Ottawa Scale.**

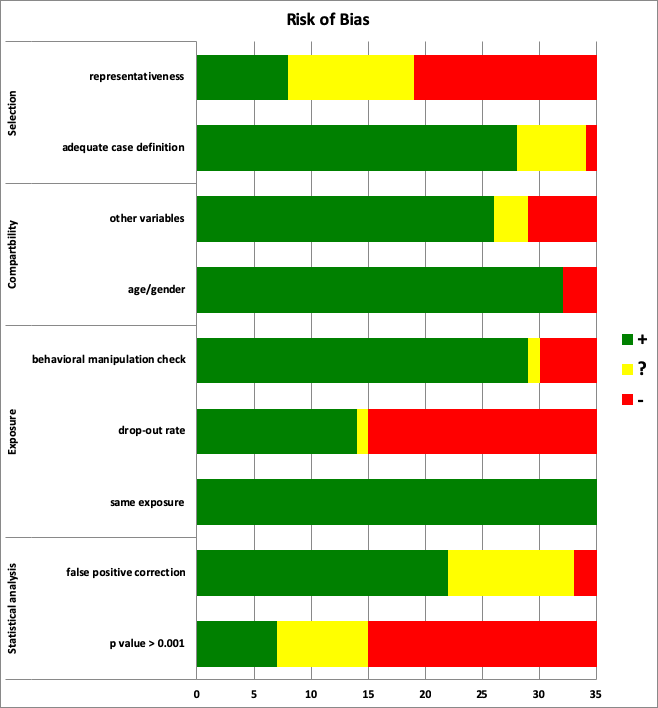

Green indicates low (RoB), yellow intermediate, and red high risk of bias for each assessment domain.

**Figure S2: Significant clusters of convergence within HC and ASD groups during face perception tasks**

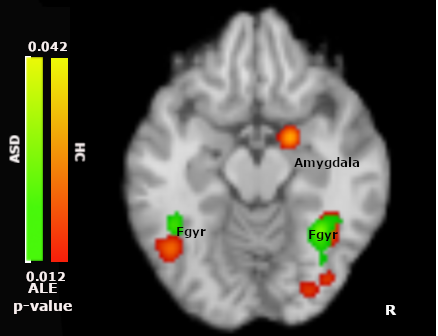

Cluster of convergence in HC group (red) and ASD group (green) during face perception tasks (p < 0.01 corrected). Amy, amygdala; FGyr, Fusiform Gyrus; ALE p-value, Activation likelihood estimation probability.

**Figure S4: Sensitivity analysis: pure face perception. Significant clusters for the comparison between autism spectrum disorder (ASD) and healthy controls (HC) (p < 0.01 corrected).**

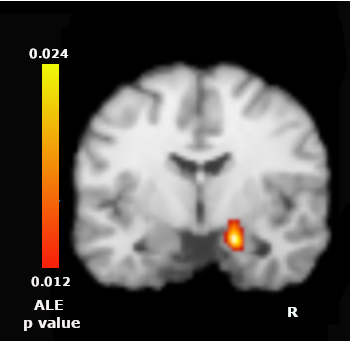

Red colors indicate higher convergence for HC (p <0.01 corrected). ALE prob, Activation likelihood estimation probability

**Figure S5: Sensitivity analysis: pooled analysis. Significant clusters considering coordinates for the ASD > HC and for the HC > ASD pooled together (p < 0.01 corrected).**

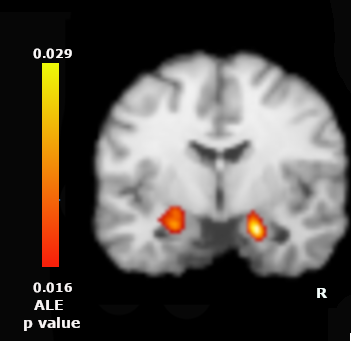

ALE prob, Activation likelihood estimation probability.

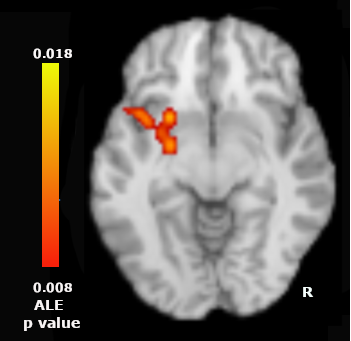
**Figure S6: Significant clusters for the comparison between autism spectrum disorder (ASD) and healthy controls (HC) in studies reporting single group activation using the convergence of difference method (p < 0.01 corrected).**

Red colors indicate higher convergence for HC p <0.01 corrected. ALE prob, Activation likelihood estimation probability

**Supplementary References**

1. Gentili C, Messerotti Benvenuti S, Lettieri G, Costa C, Cecchetti L (2018): ROI and phobias: The effect of ROI approach on an ALE meta‐analysis of specific phobias. *Human Brain Mapping*. doi: 10.1002/hbm.24492.

2. Eklund A, Nichols TE, Knutsson H (2016): Cluster failure: Why fMRI inferences for spatial extent have inflated false-positive rates. *Proceedings of the National Academy of Sciences*. 113: 7900–7905.

3. Cox RW, Chen G, Glen DR, Reynolds RC, Taylor PA (2017): fMRI clustering and false-positive rates. *Proc Natl Acad Sci USA*. 114: E3370–E3371.

4. Muller VI, Cieslik EC, Serbanescu I, Laird AR, Fox PT, Eickhoff SB (2017): Altered Brain Activity in Unipolar Depression Revisited: Meta-analyses of Neuroimaging Studies. *JAMA Psychiatry*. 74: 47–55.

5. Baron-Cohen S, Ring HA, Wheelwright S, Bullmore ET, Brammer MJ, Simmons A, Williams SCR (1999): Social intelligence in the normal and autistic brain: an fMRI study: *Social intelligence in the normal and autistic brain*. *European Journal of Neuroscience*. 11: 1891–1898.

6. Bolte S, Ciaramidaro A, Schlitt S, Hainz D, Kliemann D, Beyer A, *et al.* (2015): Training-induced plasticity of the social brain in autism spectrum disorder. *Br J Psychiatry*. 207: 149–57.

7. Brandenburg-Goddard MN, van Rijn S, Rombouts SARB, Veer IM, Swaab H (2014): A comparison of neural correlates underlying social cognition in Klinefelter syndrome and autism. *Social Cognitive and Affective Neuroscience*. 9: 1926–1933.

8. Ciaramidaro A, Bölte S, Schlitt S, Hainz D, Poustka F, Weber B, *et al.* (2018): Transdiagnostic deviant facial recognition for implicit negative emotion in autism and schizophrenia. *European Neuropsychopharmacology*. 28: 264–275.

9. Corbett BA, Carmean V, Ravizza S, Wendelken C, Henry ML, Carter C, Rivera SM (2009): A functional and structural study of emotion and face processing in children with autism. *Psychiatry Research: Neuroimaging*. 173: 196–205.

10. Critchley HD, Daly EM, Bullmore ET, Williams SCR, Van Amelsvoort T, Robertson DM, *et al.* (2000): The functional neuroanatomy of social behaviour. *Brain*. 123: 2203–2212.

11. Dalton KM, Nacewicz BM, Johnstone T, Schaefer HS, Gernsbacher MA, Goldsmith HH, *et al.* (2005): Gaze fixation and the neural circuitry of face processing in autism. *Nat Neurosci*. 8: 519–26.

12. Dapretto M, Davies MS, Pfeifer JH, Scott AA, Sigman M, Bookheimer SY, Iacoboni M (2006): Understanding emotions in others: mirror neuron dysfunction in children with autism spectrum disorders. *Nature Neuroscience*. 9: 28–30.

13. Davies MS, Dapretto M, Sigman M, Sepeta L, Bookheimer SY (2011): Neural bases of gaze and emotion processing in children with autism spectrum disorders: Neural Basis of Gaze Processing in Children with Autism. *Brain and Behavior*. 1: 1–11.

14. Deeley Q, Daly EM, Surguladze S, Page L, Toal F, Robertson D, *et al.* (2007): An event related functional magnetic resonance imaging study of facial emotion processing in asperger syndrome. *Biological Psychiatry*. 62: 207–217.

15. Doyle-Thomas KAR, Goldberg J, Szatmari P, Hall GBC (2013): Neurofunctional Underpinnings of Audiovisual Emotion Processing in Teens with Autism Spectrum Disorders. *Frontiers in Psychiatry*. 4. doi: 10.3389/fpsyt.2013.00048.

16. Duerden EG, Taylor MJ, Soorya LV, Wang T, Fan J, Anagnostou E (2013): Neural correlates of inhibition of socially relevant stimuli in adults with autism spectrum disorder. *Brain Research*. 1533: 80–90.

17. Greimel E, Schulte-Ruther M, Kircher T, Kamp-Becker I, Remschmidt H, Fink GR, *et al.* (2010): Neural mechanisms of empathy in adolescents with autism spectrum disorder and their fathers. *NeuroImage*. 49: 1055–65.

18. Greimel E, Nehrkorn B, Fink GR, Kukolja J, Kohls G, Müller K, *et al.* (2012): Neural mechanisms of encoding social and non-social context information in autism spectrum disorder. *Neuropsychologia*. 50: 3440–3449.

19. Hadjikhani N, Zürcher NR, Rogier O, Hippolyte L, Lemonnier E, Ruest T, *et al.* (2014): Emotional contagion for pain is intact in autism spectrum disorders. *Translational Psychiatry*. 4: e343–e343.

20. Herrington JD, Riley ME, Grupe DW, Schultz RT (2015): Successful Face Recognition is Associated with Increased Prefrontal Cortex Activation in Autism Spectrum Disorder. *Journal of Autism and Developmental Disorders*. 45: 902–910.

21. Holt RJ, Chura LR, Lai M-C, Suckling J, von dem Hagen E, Calder AJ, *et al.* (2014): ‘Reading the Mind in the Eyes’: an fMRI study of adolescents with autism and their siblings. *Psychological Medicine*. 44: 3215–3227.

22. Ishitobi M, Kosaka H, Omori M, Matsumura Y, Munesue T, Mizukami K, *et al.* (2011): Differential amygdala response to lower face in patients with autistic spectrum disorders: An fMRI study. *Research in Autism Spectrum Disorders*. 5: 910–919.

23. Kim S-Y, Choi U-S, Park S-Y, Oh S-H, Yoon H-W, Koh Y-J, *et al.* (2015): Abnormal Activation of the Social Brain Network in Children with Autism Spectrum Disorder: An fMRI Study. *Psychiatry Investigation*. 12: 37.

24. Klapwijk ET, Aghajani M, Colins OF, Marijnissen GM, Popma A, van Lang NDJ, *et al.* (2016): Different brain responses during empathy in autism spectrum disorders versus conduct disorder and callous-unemotional traits. *Journal of Child Psychology and Psychiatry*. 57: 737–747.

25. Koshino H, Kana RK, Keller TA, Cherkassky VL, Minshew NJ, Just MA (2008): fMRI Investigation of Working Memory for Faces in Autism: Visual Coding and Underconnectivity with Frontal Areas. *Cerebral Cortex*. 18: 289–300.

26. Lassalle A, Åsberg Johnels J, Zürcher NR, Hippolyte L, Billstedt E, Ward N, *et al.* (2017): Hypersensitivity to low intensity fearful faces in autism when fixation is constrained to the eyes: Hypersensitivity to Fearful Faces in ASD. *Human Brain Mapping*. 38: 5943–5957.

27. Loveland KA, Steinberg JL, Pearson DA, Mansour R, Reddoch S (n.d.): Judgments of Auditory—Visual Affective Congruence in Adolescents with and without Autism: A Pilot Study of a New Task Using fMRI. 19.

28. Morita T, Kosaka H, Saito DN, Ishitobi M, Munesue T, Itakura S, *et al.* (2012): Emotional responses associated with self-face processing in individuals with autism spectrum disorders: An fMRI study. *Social Neuroscience*. 7: 223–239.

29. Perlman SB, Hudac CM, Pegors T, Minshew NJ, Pelphrey KA (2011): Experimental manipulation of face-evoked activity in the fusiform gyrus of individuals with autism. *Soc Neurosci*. 6: 22–30.

30. Sabatino A, Rittenberg A, Sasson NJ, Turner-Brown L, Bodfish JW, Dichter GS (2013): Functional neuroimaging of social and nonsocial cognitive control in autism. *Journal of Autism and Developmental Disorders*. 43: 2903–2913.

31. Scherf KS, Elbich D, Minshew N, Behrmann M (2015): Individual differences in symptom severity and behavior predict neural activation during face processing in adolescents with autism. *NeuroImage: Clinical*. 7: 53–67.

32. Shafritz KM, Bregman JD, Ikuta T, Szeszko PR (2015): Neural systems mediating decision-making and response inhibition for social and nonsocial stimuli in autism. *Progress in Neuro-Psychopharmacology and Biological Psychiatry*. 60: 112–120.

33. Stanfield AC, Philip RCM, Whalley H, Romaniuk L, Hall J, Johnstone EC, Lawrie SM (2017): Dissociation of Brain Activation in Autism and Schizotypal Personality Disorder During Social Judgments. *Schizophrenia Bulletin*. 43: 1220–1228.

34. Rahko JS, Paakki J-J, Starck TH, Nikkinen J, Pauls DL, Kätsyri JV, *et al.* (2012): Valence Scaling of Dynamic Facial Expressions is Altered in High-Functioning Subjects with Autism Spectrum Disorders: an fMRI Study. *Journal of Autism and Developmental Disorders*. 42: 1011–1024.

35. Velasquez F, Qin XA, Reilly MA, Neuhaus E, Estes A, Aylward E, Kleinhans NM (2017): Neural correlates of emotional inhibitory control in autism spectrum disorders. *Research in Developmental Disabilities*. 64: 64–77.

36. Weng S-J, Carrasco M, Swartz JR, Wiggins JL, Kurapati N, Liberzon I, *et al.* (2011): Neural activation to emotional faces in adolescents with autism spectrum disorders: Neural function in adolescents with ASD. *Journal of Child Psychology and Psychiatry*. 52: 296–305.

37. Whyte EM, Behrmann M, Minshew NJ, Garcia NV, Scherf KS (2016): Animal, but not human, faces engage the distributed face network in adolescents with autism. *Developmental Science*. 19: 306–317.

38. Zürcher NR, Donnelly N, Rogier O, Russo B, Hippolyte L, Hadwin J, *et al.* (2013): It’s All in the Eyes: Subcortical and Cortical Activation during Grotesqueness Perception in Autism. (C. Schmitz, editor) *PLoS ONE*. 8: e54313.

39. Zürcher NR, Rogier O, Boshyan J, Hippolyte L, Russo B, Gillberg N, *et al.* (2013): Perception of Social Cues of Danger in Autism Spectrum Disorders. (A. Senju, editor) *PLoS ONE*. 8: e81206.
